# Characterization of Phytoplankton-Excreted Metabolites Mediating Carbon Flux through the Surface Ocean

**DOI:** 10.1101/2025.11.04.686593

**Authors:** Yuting Zhu, Hanna S. Anderson, Eli Salcedo, Samuel E. Miller, Krista Longnecker, Melissa C. Kido Soule, Sheean T. Haley, Gretchen J. Swarr, Rogier Braakman, Sonya T. Dyhrman, Elizabeth B. Kujawinski

## Abstract

The marine labile dissolved organic carbon (DOC) pool is a dynamic reservoir of thousands of molecules that cycles approximately one-quarter of Earth’s primary production within days to weeks. After excretion by phytoplankton and other microbes, metabolites are rapidly consumed, resulting in low standing concentrations (picomolar to low nanomolar). Despite the decades-long search for labile DOC sources and molecular identities, marine phytoplankton exometabolomes are not well characterized, largely due to difficulties in measuring small polar molecules in saline water. Here, we profiled the exometabolomes of six axenic phytoplankton species representing key functional groups including a diatom (*Thalassiosira pseudonana* CCMP1335), a picoeukaryote (*Micromonas commoda* RCC299), a coccolithophore (*Gephyrocapsa huxleyi* CCMP371), a diazotrophic cyanobacterium (*Crocosphaera watsonii* WH8501), and two picocyanobacteria (*Prochlorococcus marinus* MIT 9301 and *Synechococcus* WH8102). From these cultures, we quantified 56 amine- and alcohol-containing exometabolites representing 11 compound classes which in sum comprised up to 23.4% of phytoplankton-excreted DOC. We estimated that these phytoplankton-derived exometabolites could supply up to 5% of the daily carbon quota of the dominant heterotrophic bacterium SAR11 in the surface ocean. Substantial variations in exometabolite identity and concentration across phytoplankton taxa underscore taxonomic diversity as a key driver in the supply and composition of labile DOC. This taxonomic variation predicts geographic and seasonal differences in the distribution of marine dissolved metabolites that underpin the cycling of labile DOC back to CO_2_. Overall, our work suggests that phytoplankton exometabolites are key chemical currencies that mediate significant carbon fluxes within the ocean’s carbon cycle.

**Significance Statement:** Phytoplankton exometabolites are key components of the marine labile dissolved organic carbon (DOC) pool, which drives major a fraction of the oceanic carbon flux. Yet, their composition and flux are poorly constrained. Leveraging new methods, we quantified amine- and alcohol-containing exometabolites in diverse phytoplankton and found they varied taxonomically. These exometabolites accounted for up to 23.4% of excreted DOC, potentially supporting a sizable fraction of the global heterotrophic growth. Integrating our results with ecological models suggest that exometabolite composition varies geographically and seasonally in response to changing phytoplankton community structures. Our findings illuminate the long-standing “black box” of labile DOC and link taxonomic diversity to the chemical currencies underpinning the microbe-metabolite networks at the heart of the marine carbon cycle.

## Introduction

Marine phytoplankton fix ∼50 Pg carbon (C) per year (1, 2), accounting for about half of the primary production on Earth (3). Nearly half of phytoplankton-derived C (∼23 Pg C y^-1^) cycles through the labile dissolved organic carbon (DOC) pool (2, 4, 5) and is remineralized to atmospheric carbon dioxide (CO_2_) within days to weeks (6). Despite being a relatively small fraction of the total DOC pool (∼0.2 Pg C), marine labile DOC plays a major role in biogeochemical cycling, affecting the health of marine ecosystems and supporting the function of marine food webs (7–9). Phytoplankton exometabolites (i.e., biological molecules excreted into the surrounding seawater) make up a crucial fraction of marine labile DOC and facilitate inter-organism interactions within microbial communities (10, 11). Exometabolites play multiple roles in marine ecosystems, including as (a) substrates that sustain biomass production and element cycling, (b) facilitators of biochemical reactions, such as cofactors, which can be reused and exchanged among microbes, (c) ecological signals that alter microbial phenotypes, and (d) osmolytes (i.e., compatible solutes) which regulate cellular osmotic pressure (6, 12). Characterizing the specific compounds that make up phytoplankton exometabolite pools, as well as the controls on their distribution, is key to understanding the ocean carbon cycle, both today and in the context of a changing Earth system.

Phytoplankton groups occupy distinct ecological niches and express different metabolic pathways which can differ across distinct physiological and biochemical contexts, suggesting that diverse taxa may produce divergent exometabolite profiles (13, 14). A recent survey of 42 marine microbial taxa (15) suggested that presence-absence patterns of intracellular (endo-) metabolites broadly follows taxonomy, but the degree to which this is true for exometabolites is not clear. While broad qualitative (i.e., untargeted metabolomics) studies have shown that the composition of phytoplankton exometabolites generally follows taxonomy (16), specific quantitative (i.e., targeted) exometabolomics remains unexplored, constraining our knowledge of the taxonomic influences on exometabolite dynamics. Further, the distribution patterns of endo- and exometabolites within taxa are uncoupled where they have been examined. That is, *Prochlorococcus* ecotypes have been found to produce dissimilar suites of endo- and exometabolites (17), likely due to different regulation dynamics between the two pools, further highlighting the need to directly measure exometabolites. In biogeochemical models, in turn, both the pool of phytoplankton-derived DOC and its degradation by heterotrophic microbes have long been treated as ‘black boxes’ (6, 18) due to a limited understanding of the molecular-level linkages between microbial community composition, exometabolite production, and inter-organism exchange. Unraveling exometabolite chemotaxonomic patterns is thus also critical to developing models that describe key features of microbe-chemical interactions in marine waters and for elaborating the role of DOC components in macroscale ecosystem processes such as carbon cycling.

Quantification of marine exometabolites is technically challenging due to near-vanishing concentrations and short residence times, as well as interference from salts that are higher in concentration by several orders of magnitude (6). Recent advances in analytical chemistry, particularly mass spectrometry (MS) and nuclear magnetic resonance (NMR) spectroscopy, show promise for addressing these long-standing challenges (11, 19–21). For example, direct injection coupled with isocratic liquid chromatography mass spectrometry (LC-MS) can achieve sub-micromolar level analyses of exometabolites in saline culture media (22). Lower detection limits, at picomolar to nanomolar levels, were obtained with the applications of solid phase extraction (SPE) prior to targeted metabolomics analyses using LC-MS (19, 23, 24). More recently, it has been shown that chemical derivatization coupled with LC-MS based metabolomic analyses enables effective exometabolite profiling in both seawater and culture samples by targeting specific chemical functionalities (25, 26). For amine- and alcohol-containing polar metabolites, benzoyl chloride (BC) derivatization improves extraction efficiency, enhances chromatographic separation, and optimizes quantification with analyte-paired internal standards generated using BC labeled with stable isotopes (26). Here, we used BC derivatization (26) to quantify a suite of amine- and alcohol-containing exometabolites, many of which were previously inaccessible to LC-MS based metabolomics. We analyzed the exometabolomes of six taxonomically-diverse species from different phytoplankton functional groups to establish patterns of net exometabolite excretion, to understand how they varied between taxa, and to model exometabolite distribution in the surface ocean.

## Results and Discussion

### Amine- and alcohol-containing exometabolites are excreted by diverse phytoplankton functional types

Endometabolomes are known to vary between phytoplankton functional types (15, 27), but the extent to which these endometabolites are excreted to form the exometabolite pool cycled through surface ocean DOC is poorly constrained. We used a BC derivatization method (26) to determine the exometabolites excreted by key phytoplankton functional types, including a diatom (*Thalassiosira pseudonana* CCMP1335), a picoeukaryote (*Micromonas commoda* RCC299), a coccolithophore (*Gephyrocapsa huxleyi* CCMP371), a diazotrophic cyanobacterium (*Crocosphaera watsonii* WH8501), and two picocyanobacteria (*Prochlorococcus marinus* MIT 9301 and *Synechococcus* WH8102). Here, “excretion” represents *net* exudation, i.e., the sum of passive leakage and/or active exudation minus any re-uptake of exometabolites (28, 29). A total of 56 amine- and alcohol-containing exometabolites representing 11 compound classes were excreted by these taxa and were detected in the culture filtrates (Table S1). These exometabolites constituted different proportions of the excreted DOC, ranging from less than 5% of the total DOC excreted by *G. huxleyi* to nearly 24% of the total DOC excreted by the diatom *T. pseudonana* (Figure 1). With the BC method, the exometabolites were dominated by amino acids but also included diverse metabolite compound classes such as amino sugar acids, chitin-related compounds, nucleosides, nucleotides, peptides, phosphoesters, polyamines, sulfonates, vitamins, and vitamin precursors (Figure 1).

**Figure 1.**
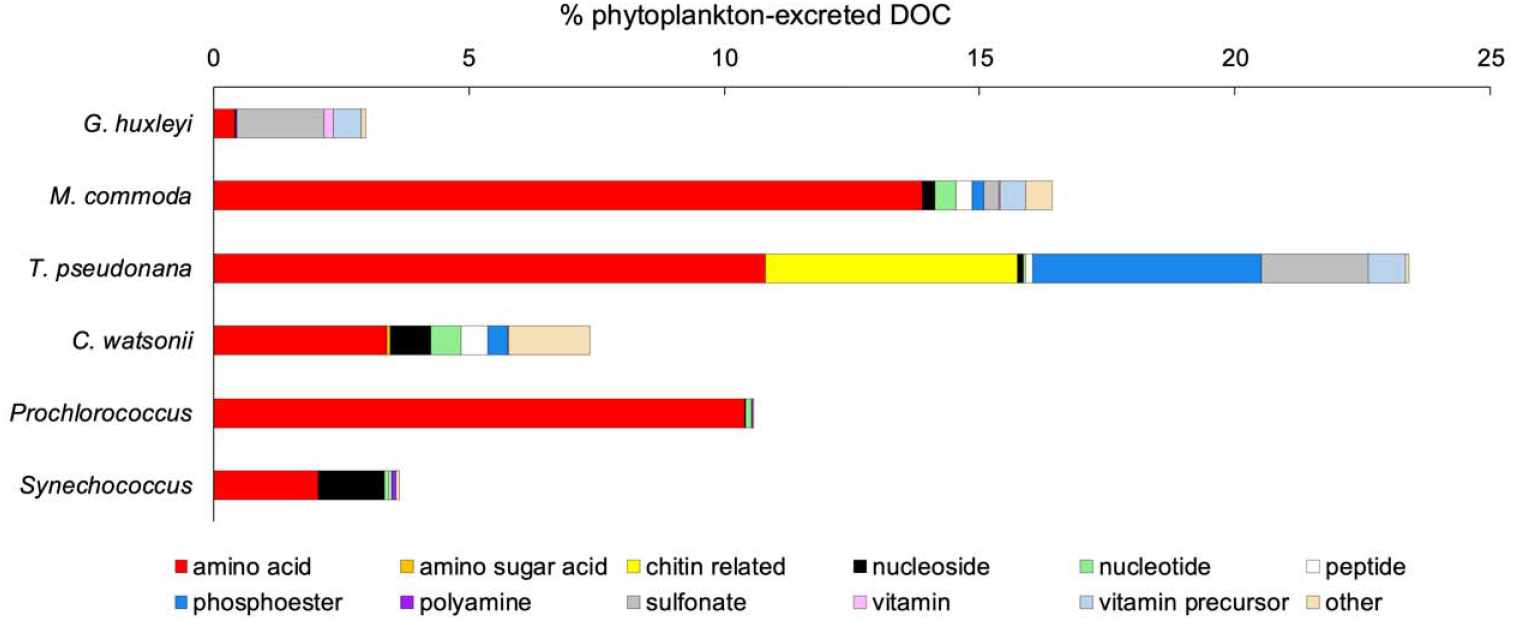
Percentage of phytoplankton-excreted DOC contributed by each exometabolite category identified for *G. huxleyi, M. commoda, T. pseudonana, C. watsonii, Prochlorococcus*, and *Synechococcus*. Exometabolites’ contributions to phytoplankton-excreted DOC were calculated by dividing the exometabolite carbon concentrations (nmol C L^-1^) by the corresponding phytoplankton species’ blank-corrected DOC (*µ*mol L^-1^) (Figure S5).

The amine- and alcohol-containing compound classes detected here, including amino acids and sulfonates, are known substrates for heterotrophic bacterial production (11) and the respiration of labile DOC back to CO_2_. To evaluate the potential scale of these processes within the carbon cycle, we estimated how much heterotrophic growth could be supported at observed excretion levels, both for total excreted DOC and for amine- and alcohol-containing exometabolites. To model heterotrophic growth, we used the widespread marine alpha-proteobacterium, *Pelagibacter ubique*, or SAR11, which reaches abundances as high as 50% of all cells in the euphotic zone (30), making it a useful reference point. Assuming that all exometabolites are bioavailable and that all are consumed with the same bacterial growth efficiency (BGE, 54%) (2, 31), we calculated that the total DOC excreted per cell per day by the different phytoplankton species could support growth of 0.09 to 34 SAR11 cells per day (Figure 2). The number of SAR11 cells that can be supported by the amine- and alcohol-containing exometabolites alone ranged from 0.01 to 3 per day (Figure 2).

**Figure 2.**
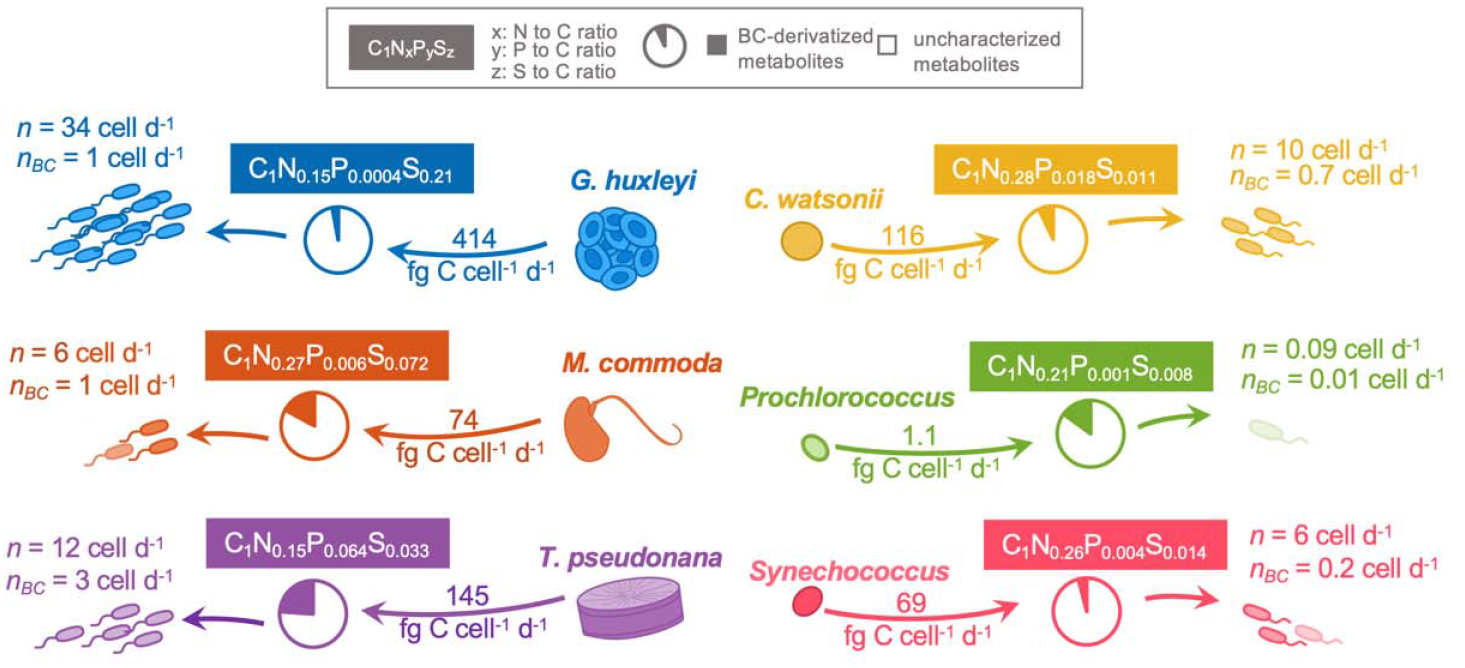
The per-cell ability of each strain to excrete fg of DOC per day in axenic cultures grown in the laboratory, the fraction of this excreted DOC made up of BC-derivatized metabolites (illustrated in pie charts), the C:N:P:S molar ratio of BC-derivatized metabolites (illustrated in colored text boxes), the number of bacterial cells that can be produced from the total excreted DOC excreted by each phytoplankton cell per day (*n*), and the number of bacterial cells that can be produced from the BC-derivatized metabolites excreted by each phytoplankton cell per day (*n*_*BC*_). We used SAR11 as a model for the bacterial cellular growth calculation, assuming 6.5 fg C cell^-1^ biomass (36) and 54% BGE (2, 31).

Larger eukaryotic cells (e.g., diatoms) excreted higher levels of DOC, allowing them to support greater heterotrophic growth on a per-cell basis. However, small prokaryotic cells (e.g., *Prochlorococcus* and *Synecococcus*) are more abundant in the global ocean and therefore likely facilitate the growth of considerable fractions of bacterial cells through DOC excretion. Hence, to further contextualize the biogeochemical scale of observed phytoplankton excretions, we estimated the levels of SAR11 growth it could sustain at a global scale using the biomass output of the MIT Darwin model (32–34). Our results suggest that, based on measured excretion levels (Figure 2) the eukaryotic phytoplankton (coccolithophores, picoeukaryotes, and diatoms < 15*µ*m) could support growth of up to 1.2–3.0×10^4^ SAR11 cells in 1 mL of seawater per day, while prokaryotic classes (diazotrophs, *Prochlorococcus*, and *Synechococcus*) could support up to 0.3– 3.3×10^4^ cells mL^-1^ day^-1^ (Figure S1). These upper limit estimates correspond to the 95^th^ percentile values of the computed SAR11 production in the model (Figure S1). Thus, despite having lower excretion levels on a per-cell basis than eukaryotes, prokaryotic phytoplankton support broadly similar levels of heterotrophic growth due to their large population sizes. Further, by converting the SAR11 biomass production reported in the Atlantic Ocean (35) assuming 6.5 fg C cell^-1^ (36), we estimated production rates of 6.8×10^4^ to 4.8×10^5^ cells mL^-1^ day^-1^, which is congruent with our calculations. For the measured amine- and alcohol-containing exometabolites, 95^th^ percentile values of SAR11 production were calculated to be on the order of 0.3–5×10^3^ cells mL^-1^ day^-1^, which corresponds to approximately 0.3–5% of total SAR11 production, assuming a typical growth rate of 1×10^5^ cells mL^-1^ day^-1^. These estimates underscore the ecological significance of these exometabolites from these representative phytoplankton taxa as key C sources supporting a substantial fraction of the ocean’s most abundant heterotrophic bacteria.

The exometabolites identified here contained C, N, P, and S at ratios that varied across the six taxa (Figure 2), reflecting how phytoplankton community composition may lead to variation in the local or regional cycling of key resource-containing exometabolites. For example, the C:S ratio (1:0.21) of the amine- and alcohol-containing exometabolites excreted by *G. huxleyi* was the highest among the six taxa and can be attributed to the dominance of sulfonates in its exometabolome (Figure 2). Similarly, phosphoesters were responsible for the relatively high C:P ratio (1:0.064) in the *T. pseudonana* exometabolome (Figure 2). In sum, amine- and alcohol-containing exometabolites can account for nearly one-quarter of total labile DOC production by certain key phytoplankton and the cycling of these compounds could support sizable populations of heterotrophic bacteria. Taxonomically-linked C:N:P:S ratios of exometabolites may vary across regions of the surface ocean depending on phytoplankton community composition, and could likely act as a control on heterotrophic production, affecting elemental fluxes and shaping ecological processes.

### Phytoplankton exometabolite profiles vary taxonomically

Clear variations were observed in the relative abundance of different compound classes among phytoplankton taxa (Figure 1). For example, the excretion of sulfonates from *G. huxleyi* and *T. pseudonana* was higher than from the other phytoplankton taxa (Figure 1), while the release of nucleosides was higher in *Synechococcus* and *C. watsonii*. The observed amino acid contributions to phytoplankton-excreted DOC were similar (10.4–13.9%) for *T. pseudonana, M. commoda*, and *Prochlorococcus* (Figure 1). While levels were somewhat lower in *Synechococcus* (2.1%) and *C. watsonii* (3.4%), the amino acid levels across most of our phytoplankton were similar to fractions (∼ 4–15%) previously reported for eukaryotic phytoplankton (37). The *G. huxleyi* exometabolome was a notable outlier, containing an amino acid pool of only 0.4% (Figure 1).

To understand the relative importance of taxonomy and possible strain-specific adaptations in shaping observed differences in excretion profiles, we quantified similarities of exometabolome profiles using hierarchical clustering analysis. There was distinct clustering of eukaryotic and the prokaryotic exometabolomes (Figure 3), with 17 exometabolites shared by the prokaryotes (Figure S2) and 13 exometabolites shared by the eukaryotes and only 5 of these shared by all six taxa across both groups. Among these shared exometabolites, 4-amino-5-aminomethyl-2-methylpyrimidine (amMP) was the only metabolite excreted by all three eukaryotic strains and absent from all prokaryotic exometabolomes, whereas the 17 exometabolites excreted by all three prokaryotic strains were also detected in eukaryotic exometabolomes. In addition, we found that four exometabolites were unique to eukaryotes (isethionic acid, amMP, chitobiose, chitotriose) and two were unique to prokaryotes (inosine and N-acetyl-muramic acid), aligning with a previous endometabolome study that found group-specific metabolites (15). Overall, these results suggest that taxonomy plays a key role in shaping metabolite excretion, while shared exometabolites highlight conserved metabolite excretion across phylogenetic groups.

**Figure 3.**
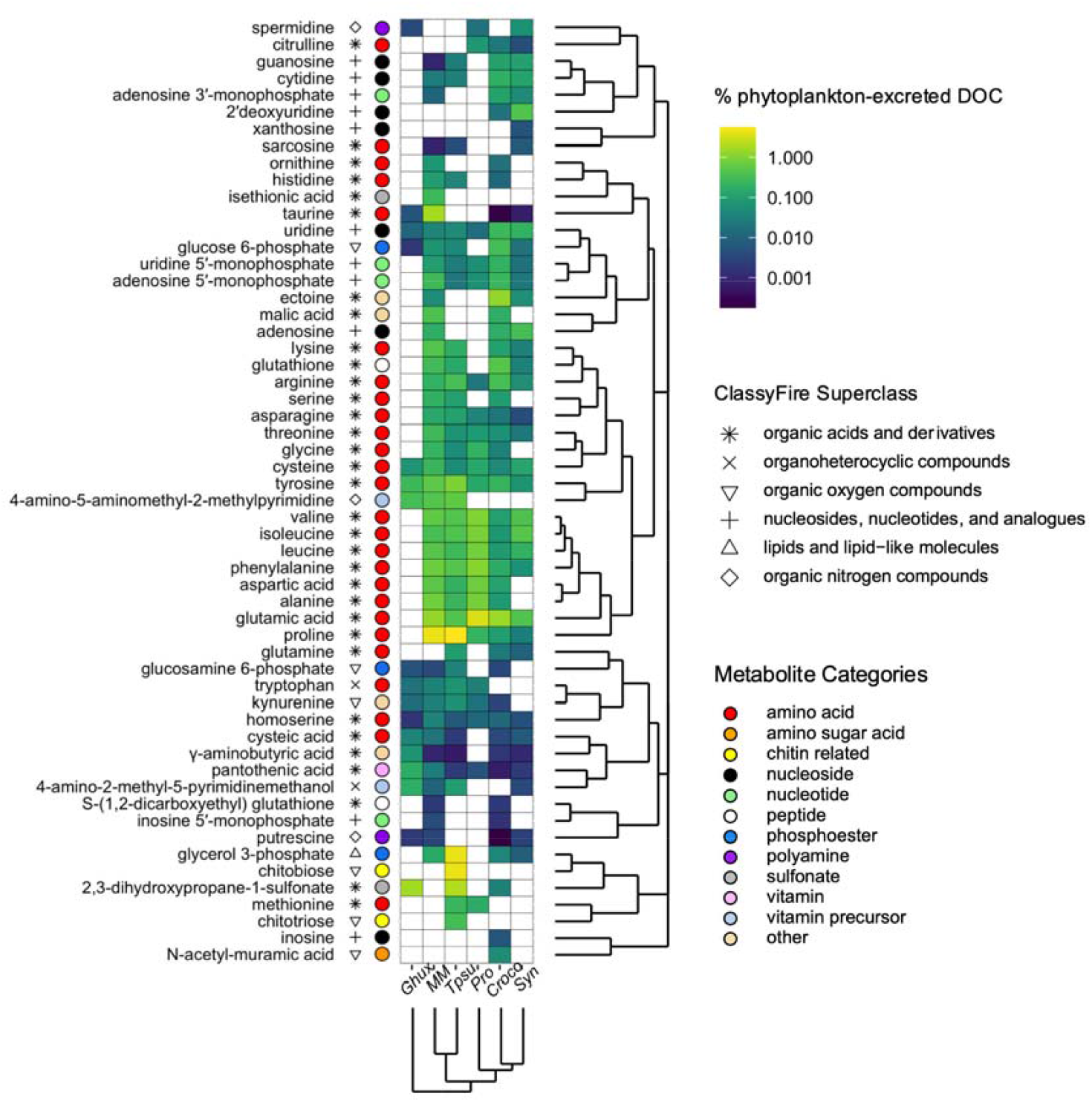
Exometabolite contributions to phytoplankton-excreted DOC in the filtrate of axenic cultures of *G. huxleyi* (*Ghux*), *M. commoda* (*MM*), *T. pseudonana* (*Tpsu*), *Prochlorococcus* (*Pro*), *C. watsonii* (*Croco*), and *Synechococcus* (*Syn*). Each identified exometabolite is paired with a symbol representing ClassyFire (59) superclass, which is a standardized chemical taxonomy. Each metabolite is also labeled with a colored marker that represents the assigned metabolite category based on biochemical functions. Exometabolites’ contributions to phytoplankton-excreted DOC were calculated by dividing the exometabolite carbon concentrations (nmol C L^-1^) by the corresponding phytoplankton species’ blank-corrected DOC (*µ*mol L^-1^). Metabolites and species were clustered based on exometabolites’ percentage contribution to phytoplankton-excreted DOC and ordered accordingly. Supplementary heatmaps are provided in Figure S6, S7, and S8 with colored boxes representing exometabolite nanomolar concentrations (nmol L^-1^), cell-specific concentrations (amol cell^-1^), and carbon concentrations (nmol C L^-1^), respectively. Cysteine is known for its instability, and so concentrations reported here are approximate. Cysteine was excluded from the most abundant metabolites discussed in the section “Distributions of high-abundance exometabolites vary with community structure in the global surface ocean”.

Several factors can potentially contribute to the observed taxonomic differentiation of phytoplankton excretion profiles, including differences in metabolic genomic potential, evolved differences in gene regulation, or variations in metabolite transport. To help differentiate among these possibilities, we examined the phytoplankton genomes or transcriptomes for the presence of relevant biosynthetic pathways. We found that 53 of the 56 exometabolites detected in our study are intermediates of biosynthetic pathways, with 41 belonging to pathways present in all six species. However, of these 41 compounds, only 4 were detected in the exometabolomes of all six species, with the remaining 37 being detected in the exometabolomes of different subsets of phytoplankton. For example, inosine and xanthosine, two purine nucleosides that are intermediates of annotated biosynthetic pathways in all six taxa, were detected as exometabolites only in *C. watsonii* and *Synechococcus*, respectively (Figure S3). N-acetyl muramic acid (a component of peptidoglycan) was present in biosynthetic pathways of all three picocyanobacteria, but was only detected in the exometabolome of *C. watsonii* (Figure S3). These results underscore how metabolite excretion is often not predictable from genomes alone and suggest that taxonomic differentiation of excretion profiles (Figure 3) is driven primarily by differences in the regulation of metabolism and/or metabolite transport rather than by differences in metabolic potential.

For some observed exometabolites we could not find an accompanying biosynthetic pathway. For example, ectoine was detected in the exometabolomes of *C. watsonii, Synechococcus*, and *M. commoda* (Figure 3), but a previously described bacterial ectoine biosynthesis pathway (38) was not present in any of the species in our study. Ectoine has previously been measured in the metabolome of an axenic diatom isolate that had only parts of a recognized ectoine biosynthetic pathway (39). Our exometabolomes thereby expand the diversity of taxa in which ectoine production has been detected in the absence of an ectABC gene cluster that encodes the canonical ectoine biosynthesis pathway (40), highlighting this as a key area of future study. Another example comes from kynurenine, an oxidized derivative of tryptophan. Kynurenine was excreted by the three eukaryotes whose genomes encode kynurenine production pathways, but also by *Prochlorococcus*, which lacks a corresponding biosynthetic pathway (Figure S3). In a previous study, kynurenine was not detected in *Prochlorococcus* MIT9301, but it was detected in the exometabolomes of strains MIT0801 and MIT9313 (17), suggesting it is a common exometabolite of this genus. It has been hypothesized (41) that marine cyanobacteria produce kynurenine non-enzymatically through oxidation by reactive oxygen species of tryptophan residues in the proteins of photosystem II (42). Together, these findings further highlight how excretion is often not predictable from genomes and demonstrate the power of integrating metabolomics with other omics data to identify missing or incomplete gene annotations (Supplementary Text) and refine key genomes of interest.

### Distributions of high-abundance exometabolites vary with community structure in the global surface ocean

For the 56 amine- and alcohol-containing exometabolites quantified here, their individual contributions to phytoplankton-excreted DOC across the six taxa ranged from 0.00018% to 5.8% (Figure 3). Combining the lists of the ten most abundant (hereafter “top 10”) exometabolites in terms of percent DOC contribution for each taxon generated a list consisting of 34 distinct exometabolites (Figure 4). Thirteen exometabolites, 9 of which were amino acids, were among the top 10 exometabolites for multiple phytoplankton species, highlighting their potential significance in labile DOC cycling in seawater (Figure 4). Collectively, the excretion of these top 10 exometabolites represents a critical flux of C currencies to the labile DOC pool which underpins interactions and carbon fate in a network of surface ocean autotrophs and heterotrophs. For example, all three eukaryotic species examined excreted thiamin (vitamin B_1_) precursors 4-amino-2-methyl-5-pyridinemethanol (HMP) and amMP (Figure 3). HMP and amMP were among the most abundant exometabolites detected for *G. huxleyi*, contributing to 0.2% and 0.4% of this strain’s total excreted DOC pool, respectively. Both observations add to the growing recognition that thiamin precursors are important currencies in marine ecosystems, where they can support the growth of a variety of microbes (43). Indeed, some strains of the highly abundant SAR11 group of bacterioplankton are known to be auxotrophic for HMP and cannot use exogenous thiamin for growth (44).

**Figure 4.**
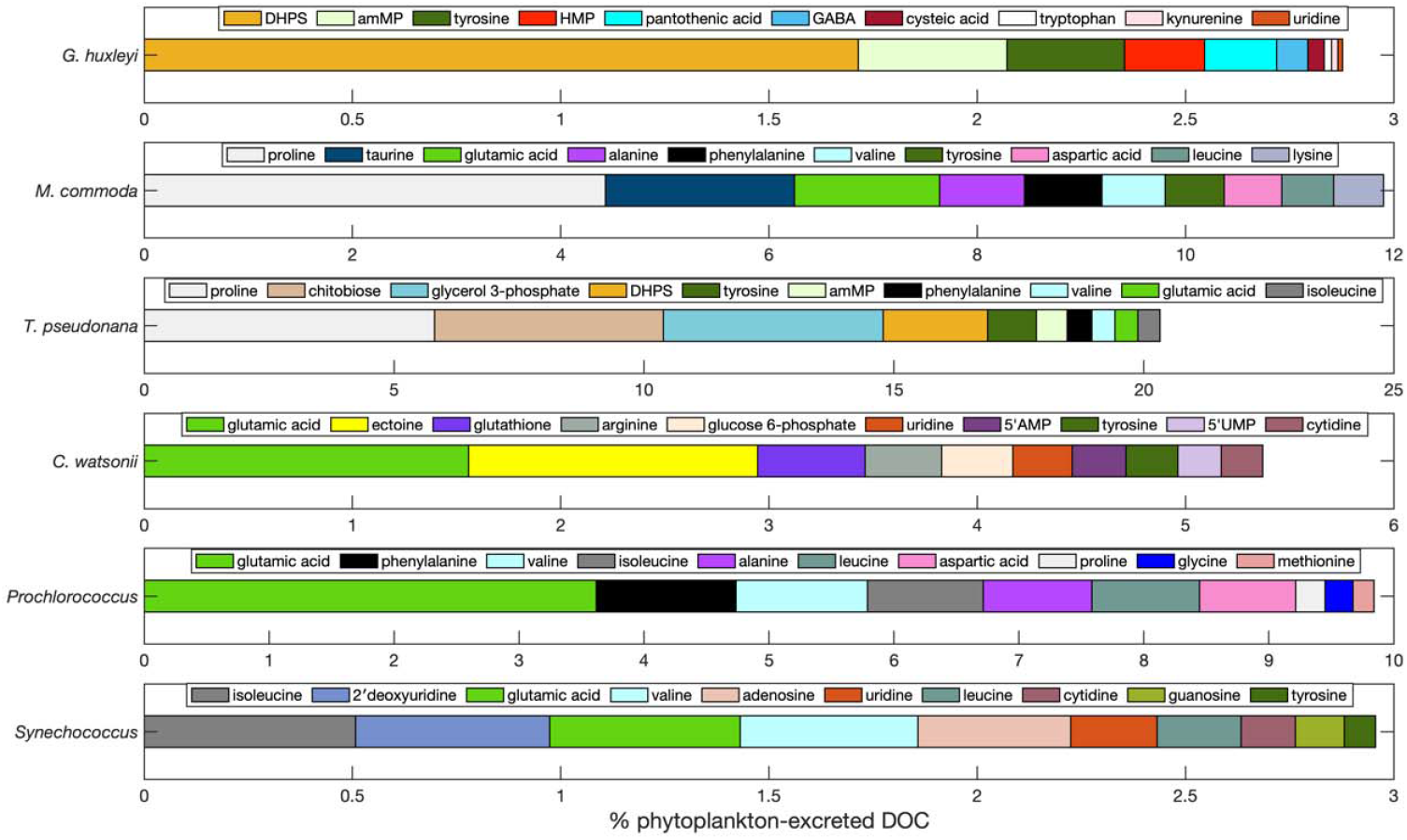
Ten exometabolites with the highest contribution to phytoplankton-excreted DOC in the filtrate of axenic cultures of *G. huxleyi, M. commoda, T. pseudonana, C. watsonii, Prochlorococcus*, and *Synechococcus*. Each exometabolite is labeled by a unique color. Note the differences in x-axis scale among the six phytoplankton species.

Many of the top 10 exometabolites have corresponding transporters characterized in heterotrophic bacteria, reinforcing their significance in the cycling of labile DOC and their roles in mediating microbial interactions. For example, glycerol 3-phosphate, a dissolved organic phosphorus compound, is the third most abundant exometabolite produced by *T. pseudonana* (4.4% of excreted DOC), consistent with a previous NMR study that noted the presence of this compound in the *T. pseudonana* exometabolome (11). Gene expression patterns for a putative glycerol 3-phosphate transporter in co-cultures of *T. pseudonana* and *Ruegeria pomeroyi*, an alpha-proteobacterium, suggested that glycerol 3-phosphate was a key C and/or P substrate supporting growth of the coastal ocean heterotroph (45). The exometabolite measurements here are consistent with this hypothesis, with the scale of glycerol 3-phosphate excreted by *T. pseudonana* suggesting it could be an important source of C and P in marine waters. *R. pomeroyi* has annotated transporters for a number of the top 10 exometabolites excreted by phytoplankton screened here, including ectoine, 2,3-dihydroxypropane-1-sulfonate (DHPS), taurine, isethionic acid, cysteic acid, malic acid and putrescine (46), and gene expression studies further suggest it may also use proline, alanine, and valine (47). The ability of the model marine heterotroph *R. pomeroyi* to use many of the metabolites we have identified as major phytoplankton exometabolites supports the idea that they are key chemical currencies in ocean ecosystems.

In the picocyanobacteria, coccolithophore, diatom, picoeukaryote, and diazotroph taxa assayed here, the exometabolites with the largest percent contribution to phytoplankton-excreted DOC were glutamic acid, DHPS, proline, and ectoine (Figure 4). Glutamic acid appears on the top 10 list for four of six phytoplankton species; it is the most abundant exometabolite for *Prochlorococcus* (3.6% of excreted DOC), and the second most abundant exometabolite for *C. watsonii* (1.6% of excreted DOC) (Figure 4). While *in situ* measurements of surface ocean amine- and alcohol-containing exometabolites are relatively rare (26), abundances of glutamic acid (up to 50 nM) were recently reported in reef waters off the southern coast of the US Virgin Islands (25). The exometabolite profiles here suggest that glutamic acid in natural seawater is linked with taxonomically diverse phytoplankton producers. DHPS is a key C and S cycle intermediary in the labile DOC pool due to its role in mediating autotroph-heterotroph interactions such as between *T. pseudonana* and *R. pomeroyi* (10, 11). Past work suggests diatoms and coccolithophores could accumulate mM levels of intracellular DHPS (48), and here we found that DHPS was one of the top 10 most abundant exometabolites of *G. huxleyi* (1.7% of excreted DOC) and *T. pseudonana* (2.1% of excreted DOC) (Figure 4). Although DHPS was previously thought to be an exclusively eukaryotic endometabolite (15), it was detected here in the excreted DOC pool of a cyanobacterium (*C. watsonii*) (Figure 3), suggesting cyanobacteria may play a larger role in organic S cycling and related microbial interactions than previously known. Last, a variety of multifunctional currencies are significant contributors to phytoplankton-excreted DOC. This includes proline, the most abundant exometabolite for both *M. commoda* (4.4% of excreted DOC) and *T. pseudonana* (5.8% of excreted DOC) (Figure 4). Proline can be assimilated by bacteria as C and energy sources (46, 47) and also serves as an osmolyte, or compatible solute, for both eukaryotes and prokaryotes (49). Similarly, ectoine, which accounts for 1.4% of *C. watsonii*’s excreted DOC (Figure 4), is a known osmolyte, helping bacteria and archaea survive extreme environments (50, 51), and can be used by marine bacteria as a growth substrate (e.g., *R. pomeroyi*) (46, 52).

To evaluate the geographic and seasonal distribution of these abundant exometabolites (glutamic acid, DHPS, proline, and ectoine) in the field, we modeled their excretion as a function of their percent contribution to DOC in six known phytoplankton functional types represented in the MIT Darwin model. These include coccolithophores, diazotrophs, *Prochlorococcus, Synechococcus*, picoeukaryotes, and diatoms < 15 *µ*m) (32–34) (Figure 5), which serve as analogs to the phytoplankton assayed here. Proline excretion in this modeled output was greatest in the high latitudes and varied seasonally with shifts in diatom populations (Figure 5). DHPS excretion had a similar pattern to proline, with diatoms driving the patterns of both exometabolites (Figure 5). Besides DHPS, proline had the highest modeled excretion of any exometabolite, reinforcing its potential importance in the cycling of labile DOC, particularly in regions with large diatom populations. DHPS and proline excretion patterns were distinct from glutamic acid, which had higher excretion in the subtropics than the diatom-dominated proline and DHPS distributions (Figure 5). Glutamic acid excretion by *Prochlorococcus* controlled the subtropical distribution of this exometabolite, which does not vary seasonally like at high latitudes (Figure 5). Ectoine was only excreted at high levels by *C. watsonii*, which confined the modeled distribution of this exometabolite to tropical and subtropical regions with abundant diazotroph populations and resulted in lower fluxes than the other modeled exometabolites (Figure 5). These data demonstrate how phytoplankton excretion patterns and community structure can shape regional distributions and seasonal cycles of the labile DOC pool. The underlying heterotrophic cross-feeding networks, which underpin major components of the labile DOC cycle, likely also vary considerably across ocean regions in conjunctions with phytoplankton group variations.

**Figure 5.**
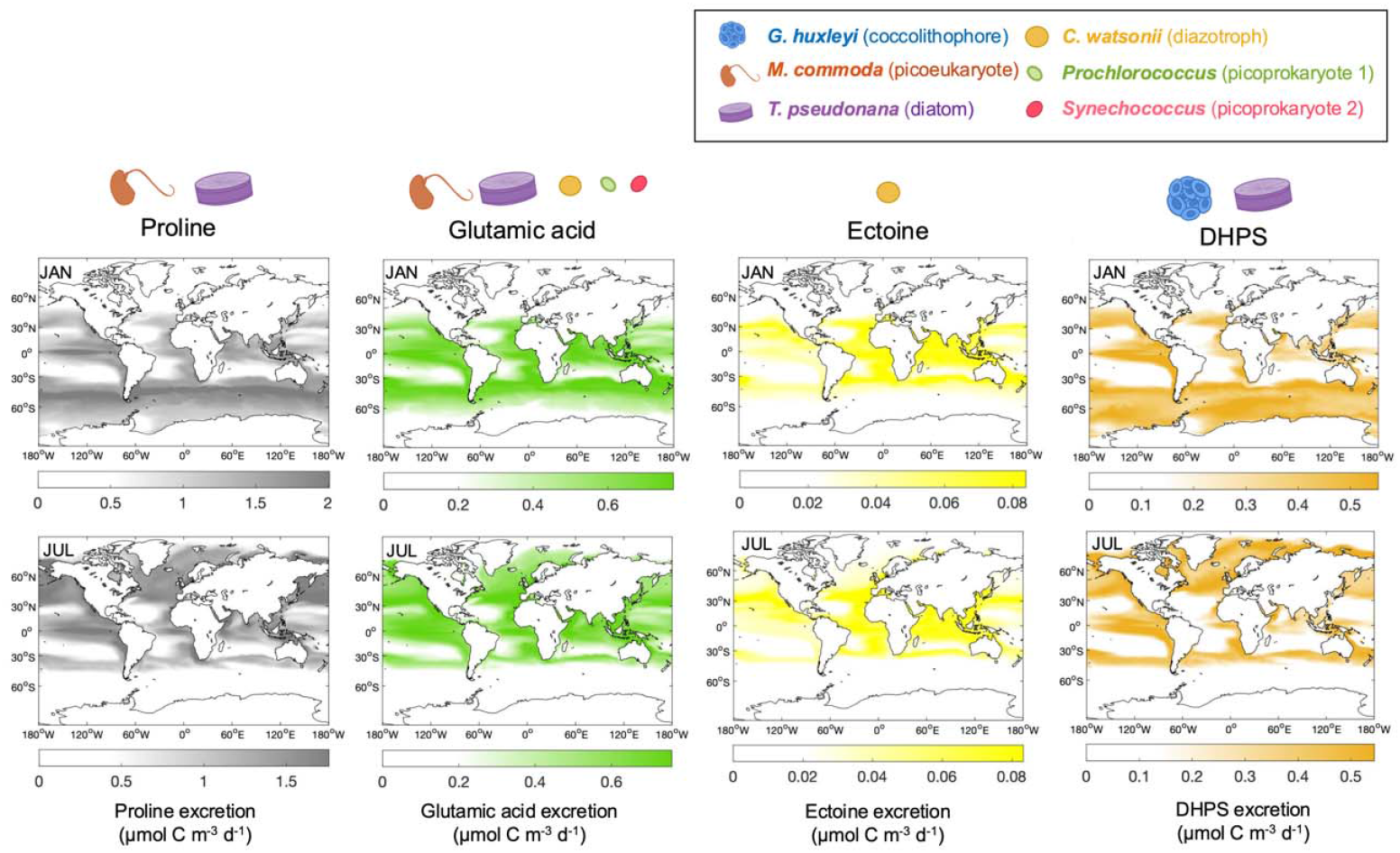
January and July maps of estimated proline, glutamic acid, ectoine and DHPS excretion in the top 100 m seawater of the global ocean. Labels indicate producers for which the metabolite is among the top 10 exometabolites (Figure 4). Surveyed phytoplankton and their functional group are indicated by color.

The quantitative determination of exometabolite concentrations in this work identified the amine- and alcohol-containing exometabolites excreted in high abundance by different phytoplankton functional groups. These exometabolites are likely key chemical currencies that shape the autotroph-heterotroph interactions which control ocean-scale carbon fluxes through marine DOC. Unraveling how phytoplankton exometabolites shape the labile DOC cycle, and thereby the return of CO_2_ to Earth’s atmosphere, is essential to understanding carbon flux, microbial-chemical networks, and ecosystem function in the context of a changing ocean.

## Materials and Methods

### Phytoplankton Culturing and Harvest

Axenic cultures included *G. huxleyi* CCMP371 (isolated from the Sargasso Sea, North Atlantic, 1987), *M. commoda* RCC299 (isolated from the South Pacific, 1998, and made axenic in 2006; synonymous with CCMP2709), *T. pseudonana* CCMP1335 (isolated from Moriches Bay, Long Island, New York, USA, 1958), *Synechococcus* sp. WH8102 (isolated from the Atlantic Ocean, 1986), *C. watsonii* WH8501 (isolated from the tropical North Atlantic Ocean, 1988), *Prochlorococcus* MIT 9301 (isolated from the North Atlantic Ocean, 1990) (Table 1). *G. huxleyi, M. commoda*, and *T. pseudonana* were grown in L1 growth media prepared according to Guillard and Hargraves (53), and *Synechococcus* was grown in SN media prepared according to Waterbury et al. (54), all with a base of 0.2-*µ*m filtered coastal seawater collected from Vineyard Sound, MA, United States. *G. huxleyi* media excluded Si. *C. watsonii* was grown in SO media, which followed the SN media recipe but omitted NaNO_3_, with a base of 75% Sargasso seawater (diluted to 75% with MilliQ). *Prochlorococcus* was grown in Pro99 in a Sargasso seawater base prepared according to Moore et al. (55). Axenic strains of *C. watsonii, G. huxleyi, M. commoda, Synechococcus*, and *T. pseudonana* were inoculated into 25 mm borosilicate glass culture tubes containing 25 mL of sterile media. Axenic *Prochlorococcus* was inoculated into borosilicate culture tubes containing 35 mL of sterile media. Nine biological replicates were monitored for all strains on a 14:10 hour light-dark cycle, with the exception of *Prochlorococcus*, which was grown on a 13:11 hour light-dark cycle. Isolates were cultured with light intensities ranging from 45–130 *µ*mol photons m^-2^ s^-1^ depending on taxonomy (Table 1).

**Table 1.**
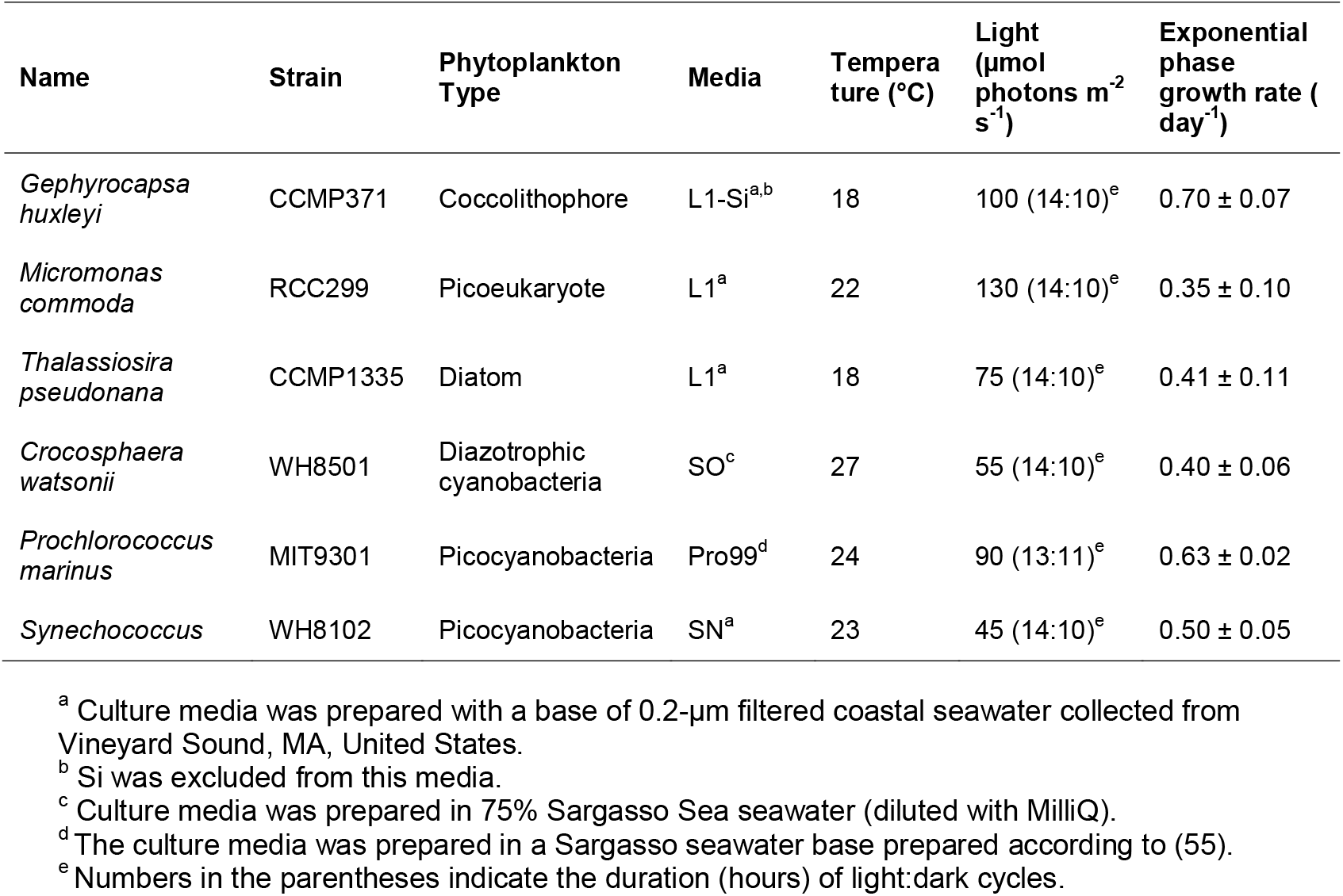
Information regarding the phytoplankton strains and growth conditions.

Growth was monitored daily by *in vivo* chlorophyll fluorescence on a Turner Designs 10-AU fluorometer. Measurements for all genera except *Prochlorococcus* were made at the same time each day, 3-4 hours after the beginning of the light cycle, to avoid diel changes in cell physiology. Daily sampling times for *Prochlorococcus* varied as cells divide only at night when acclimated to diel conditions, and thus sampling time throughout the day is less likely to influence cell counts. Six (6) replicates of each isolate were harvested by gentle vacuum filtration during exponential growth to maximize metabolite generation. The remaining 3 cultures were monitored as described above until they reached stationary phase, which occurred between 3–7 days after harvesting depending on the strain (Figure S4). For all six phytoplankton strains, cells were filtered onto filters (47 mm, 0.2 *µ*m Omnipore filters, Whatman) and the filtrates were collected into combusted (450 °C for at least 4 h) glass side-arm flasks. The vacuum pump pressure did not exceed ∼5” Hg to minimize cell lysis and endometabolite contamination in filtrate samples. Filtrates were transferred into pre-combusted 40 mL amber vials. Culture media blanks (with no algal biomass) for all media types were also filtered using the same method. All filtrate samples were stored upright at −20 °C until analysis. Growth rates during the exponential phase were calculated using relative fluorescent units (RFUs) for all strains with the exception of *Prochlorococcus*, where daily cell count data was used, and then averaged across replicates.

### Cell Counts

For all species except *Prochlorococcus*, the final cell yields were evaluated via microscopy. Briefly, 1–2 mL of well-mixed cultures were removed immediately prior to harvesting and preserved in paraformaldehyde (for *Synechococcus*; 0.24% final concentration) or neutral Lugol’s iodine solution (for all others; 2% final concentration). *Synechococcus* samples were diluted and collected on a filter (black 0.2 *µ*m, 25 mm polycarbonate filters, Whatman) for cell counts using an epifluorescence microscope (63x, oil immersion), while Lugol’s-preserved samples were counted using a hemocytometer and light microscope (10x). *Prochlorococcus* cell counts were obtained using flow cytometry. Briefly, flow cytometry samples were run live on a Guava EasyCyte flow cytometer (Cytek Biosciences) for 10,000 counts or 5 minutes each. Cells were excited with a blue 488 nm laser and cell counts, cell size, and chlorophyll content were analyzed using FlowJo.

All isolates but *Prochlorococcus* were tested to ensure they were axenic before analysis. Axenic testing consisted of adding 1 mL of each culture to 15 mL of bacterial growth media, which was stored in the dark at room temperature for up to a week and checked visually for bacterial biomass growth. Bacterial growth media was prepared by adding 5 g peptone and 10 g malt extract to 1 L filtered coastal seawater prior to autoclaving (after Freshwater Test Media, NCMA). To check that each *Prochlorococcus* sample was derived from an axenic culture, samples from each timepoint were preserved for flow cytometry done with identical specifications as described above. Briefly, 10 *µ*L of each sample was added to 990 *µ*L of 0.2 *µ*m filtered, autoclaved Sargasso seawater. 5 *µ*L of glutaraldehyde was added, and the tube was inverted and incubated for 10 minutes in the dark. Samples were flash frozen in liquid nitrogen and stored at −80°C until they could be thawed and stained with SYBR Green I for 55 minutes (Lonza). Cells were analyzed via flow cytometry for chlorophyll fluorescence, cell size, and SYBR Green I stained DNA fluorescence.

### Dissolved Organic Carbon Analysis

Five (5) mL of filtrate samples and media blanks were diluted with 25 mL of Milli-Q water, acidified to pH∼3 with concentrated hydrochloric acid, and stored at 4°C until processed on a Shimadzu TOC-L total organic carbon analyzer. Measurements were made using potassium hydrogen phthalate as a standard solution. DOC concentration was determined by subtracting the instrument blank area from the average peak area and dividing by the slope of the standard curve. Comparison with consensus reference material purchased from the Hansell lab at the University of Miami was conducted regularly to ensure the accuracy of these measurements.

### Sample Preparation and Derivatization

An 11-point standard curve was prepared in aged, 0.1-*µ*m filtered seawater collected from offshore Caribbean Sea, with standard additions of 0, 0.001. 0.01, 0.05, 0.1, 0.5, 1, 5, 10, 50, 100 ng for each metabolite. Three 6-point matrix-matched standard curves were prepared using media blanks of *T. pseudonana* (L1), *Synechococcus* (SN), and *C. watsonii* (SO), with standard additions of 0, 0.01, 0.1, 1, 10, 100 ng for each metabolite. We also prepared a matrix-matched standard curve for *Prochlorococcus* (Pro99), with standard additions of 0, 0.01, 0.1, 1, 10, 50 ng for each metabolite. All standards were prepared in 5 mL.

Standards, filtrate samples, and media blanks were derivatized following the method published in (26). The derivatization and sample preparation protocols together with pictures and video clips are available online (https://www.protocols.io/workspaces/kujawinski-lab). Briefly, each 5 mL sample was basified with 150 *µ*L of 8 M NaOH, derivatized with 1 mL BC working solution for 5 min, and then acidified with 75 *µ*L phosphoric acid. The BC working solution was prepared by mixing 95 mL acetone (Optima, ACROS Organics) with 5 mL BC (99%, ACROS Organics) and was used within 36 h after preparation. Two sets of stable isotopically labeled internal standards (SIL-IS) were prepared by derivatizing standard mixes of metabolites in seawater (aged, 0.1-*µ*m filtered seawater collected from offshore Caribbean Sea) using BC working reagent prepared with ^13^C_6_-BC (99% ^13^C, Sigma-Aldrich) or D_5_-BC (99% D, Cambridge Isotope Laboratories, Inc.), respectively. SIL-IS were derivatized following the same protocol as the samples.

Each derivatized standard, filtrate, or media blank was spiked with 498 pg addition of ^13^C_6_-labeled SIL-IS and 9.96 ng addition of D_5_-labeled SIL-IS and was dried using a vacufuge (Eppendorf) until ≥ 95% (by weight) of acetone was removed. Upon acetone removal, the liquid was transferred onto a preconditioned (6 mL of methanol followed by 24 mL of 0.01 M HCl) Bond Elut PPL cartridge (1g/6 mL, Agilent) and loaded by gravity. The samples were eluted with 6 mL of methanol by vacuum followed by evaporation to near dryness in a vacufuge. Samples were reconstituted with 100 *µ*L 5% acetonitrile in Milli-Q water, vortexed thoroughly, and then centrifuged at 12000 g, 22 °C for 5 min. The supernatant was transferred to LC vials with glass inserts which were previously spiked with 5 *µ*L of acetonitrile. The samples were stored at 4 °C until LC-MS analyses.

### UHPLC-ESI-MS/MS Operation

Targeted metabolomics analyses were performed using an ultrahigh performance liquid chromatography system (Vanquish UHPLC, Thermo Scientific) coupled to a heated electrospray ionization source (H-ESI) and an Orbitrap Fusion Lumos Tribrid mass spectrometer (Thermo Scientific) following the approach described in (26). A Waters Acquity HSS T3 column (2.1 mm × 100 mm, 1.8 *µ*m), equipped with an Acquity HSS T3 VanGuard Pre-column, was used for chromatographic separation at 40 °C. The column was eluted at 0.5 mL min^-1^ with a combination of solvents: A) 0.1% formic acid in water and B) 0.1% formic acid in acetonitrile. The chromatographic gradient was as follows: 1% B (0–0.5 min), 10% B (0.5–2.0 min), 10% B (2.0– 5.0 min), 25% B (5.0–7.0 min), 25% B (7.0–9.0 min), 50% B (9.0–12.5 min), 95% B (12.5–13.0 min), 95% B (13.0–14.5 min), 1% B (14.5–14.6 min), and 1% B (14.6–16.0 min). The autosampler was set at 4 °C. Individual autosampler injections (5 *µ*L each) were used for negative and positive ion mode analyses. The first 0.8 min of flow was diverted to waste after passing through the column. Other instrument parameters were: ESI voltages = 3600 V (positive) and 2600 V (negative); source gases = 55 (sheath), 20 (auxillary), and 1 (sweep); capillary temperature = 350°C; vaporizer temperature = 400 °C. MS data from 170–1000 m/z were collected at resolution 60,000 FWHM (at m/z 200), automatic gain control (AGC) at 4e5, and max injection time 50 msec. MS/MS data were triggered using a targeted mass list with m/z and retention time window for each derivatized analyte, and collected at resolution 7,500 FWHM, AGC 5e4, and max injection time 22 msec using higher energy collisional dissociation (HCD) with 35% collision energy and intensity threshold 2e4. Parent ions were isolated within the quadrupole at a width of 1 m/z.

### MS Data Processing

Metabolite identification and peak area integration were performed using Skyline (56, 57). MS data and retention times were used for peak identification and integration. MS/MS data were used to confirm the compound identification. Metabolite concentrations were quantified using calibration curves with peak area ratios of the non-labeled (light) metabolite added (ng) to the heavy-D_5_ or heavy-^13^C_6_ internal standard. The MATLAB script (considerSkyline.m) used for processing Skyline output is available at the Kujawinski Lab SkyMat repository (https://github.com/KujawinskiLaboratory/SkyMat). For several metabolites, calculated concentrations exceeded the 11-point calibration curve’s upper limit (e.g., 100 ng). To determine the limit of linearity (LOL), the standard curve was extended to 1500 ng. Glycerol 3-phosphate was the only metabolite with its concentrations in *T. pseudonana* filtrates (961–1108 ng) slightly exceeding the calculated LOL (1000 ng), while the concentrations of the rest of the 55 metabolites were all below the LOL. Therefore, no further correction was applied to the concentrations quantified using the 11-point calibration curve. Metabolite concentrations were filtered based on the calculated limit of detection (LOD), which was calculated using a subset standard curve taken from the lower end of the 11-point calibration curve. The subset standard curve contains a minimum of 4 data points (a blank and three non-zero points) and encompasses the limit of quantification (LOQ). LOD and LOQ were calculated as 3.3 and 10 times the standard deviation of the y-intercept divided by the slope, respectively. Metabolite concentrations lower than the calculated LODs were replaced with zeros. For most of the metabolites, these steps yielded two data tables for each ionization mode, one based on ^13^C_6_ normalization and the other based on D_5_ normalization.

A matrix correction was conducted by comparing the 11-point calibration curve prepared in seawater and the 6-point matrix-matched standard curves prepared in media blanks and normalized with the corresponding SIL-IS. Statistical differences between the slopes of the two standard curves were assessed for each matrix type, employing an analysis of the variance procedure that was introduced in (58). If the slopes were statistically different, a matrix correction factor was calculated by dividing the slope of the matrix-matched calibration curve by the slope of the calibration curve prepared in seawater. The adjusted concentration was then determined by dividing the metabolite concentration by this matrix correction factor.

To choose which SIL-IS and ion mode to use, the 95% prediction interval for the metabolite concentration at each sample point was calculated using each calibration curve. For each metabolite, the SIL-IS that yielded a standard curve with the lowest average 95% prediction interval across the range of sample concentrations was selected. For metabolites analyzed in both positive and negative ion modes, we selected the ion mode with the lowest 95% prediction interval for the selected calibration curve.

In the final data table, presence of exometabolites was determined based on whether their concentrations in filtrates are statistically higher (Student’s t-test, *P*< 0.05) than the media blanks. Each identified exometabolite is assigned with a ClassyFire (59) superclass, which is a standardized chemical taxonomy. We also assigned each metabolite with a category based on biochemical functions. Nanomolar concentrations were obtained by subtracting average concentrations in filtrate samples by average concentrations in media blanks. Carbon concentrations were obtained by multiplying blank-corrected, average nanomolar concentrations by number of carbon atoms in each molecule. Cell-specific concentrations were obtained by dividing nanomolar concentrations by the cell counts (Figure S9).

### Genome and transcriptome-based metabolite synthesis

The biosynthetic potential of the six phytoplankton strains to produce measured exometabolites was assessed from reference genomes of *C. watsonii* WH8501, *M. commoda* RCC299, *Prochlorococcus* MIT9301, *Synechococcus* WH8102, and *T. pseudonana* CCMP1335 (60–64), and a transcriptome of G. huxleyi CCMP371 (65) given its lack of a published genome. Exometabolites were first sought in the Kyoto Encyclopedia of Genes and Genomes (KEGG) (v115.1) COMPOUND database and KEGG pathways for the five reference genomes, which were present in the KEGG GENOME database with KEGG Ortholog (KO) gene annotations (66). The *G. huxleyi* transcriptome sequence assembly (GenBank accession GHJP00000000.1) was loaded as an anvi’o (v8 dev) contigs database (67). Genes in assembled transcripts were called using TransDecoder (v3.0.1) (Haas, B.J. https://github.com/TransDecoder/TransDecoder), annotated with KOs (anvi-run-kofams), and inspected in the context of KEGG pathways (anvi-draw-kegg-pathways) for exometabolite biosynthetic potential. For exometabolites not in the KEGG COMPOUND database or not identified as being substrates or products in organismal KEGG pathways, we searched for documentation of potential biosynthetic routes, including homologous proteins and nonenzymatic reactions likely to catalyze biosynthesis in cells (see Supplementary Text). Exometabolites found to be directly associated with a reaction integrated with an organismal biosynthetic pathway were included in the biosynthesizable pool of metabolites.

### Calculation of carbon flux and supported bacterial cellular growth

The per-cell ability of each phytoplankton strain to excrete fg of DOC per day in axenic cultures grown in the laboratory was calculated by dividing the total phytoplankton-excreted DOC (*µ*mol C L^-1^) (Figure S5) by the daily average concentration of new phytoplankton cells from the start of culturing until harvesting (cells mL^-1^ day^-1^). The number of new bacterial cells that can be produced from the total excreted DOC by each phytoplankton cell per day was calculated using SAR11 as a model, assuming 6.5 fg C cell^-1^ (36) and 54% bacterial growth efficiency (BGE) (2, 31).

On the global scale, the climatological biomass (mmol C m^-3^) output from the MIT Darwin model (32–34) for coccolithophore, picoeukaryote, diatom (< 15 *µ*m fractions), diazotroph, and picoprokaryote size fractions 1 (analogue for *Prochlorococcus*) and 2 (analogue for *Synechococcus*) were downloaded from Simons Collaborative Marine Atlas Project (CMAP; https://simonscmap.com/) (68) using the MATLAB client - matcmap (https://github.com/simonscmap/matcmap). The cellular density (cells m^-3^) of each phytoplankton type was computed by dividing the biomass output by the respective carbon quota. Carbon quotas of 46 fg C cell^-1^ and 213 fg C cell^-1^ were assumed for *Prochlorococcus* and *Synechococcus*, respectively, based on values reported by Bertilsson et al. (69). Coccolithophore, picoeukaryote, diatom, and diazotroph carbon quotas were estimated from cell volumes using equations given in Menden-Deuer and Lessard (70). The number of SAR11 cells supported by excreted DOC was calculated by multiplying phytoplankton cell densities by the calculated per-cell ability of each of the six phytoplankton strains examined in this work to support production of new SAR11 cells per day.

### Modeling of excretion patterns

In the top 100 m of seawater in the global ocean, the excretion of glutamic acid, proline, ectoine, and DHPS was calculated using the MIT Darwin model’s climatological biomass output for coccolithophores, picoeukaryotes, diatoms (<15 *µ*m), diazotrophs, and picoprokaryote size fractions 1 (analogous to *Prochlorococcus*) and 2 (analogous to *Synechococcus*). The cellular densities (cells m^-3^) of these phytoplankton types were computed following the procedure described above. These values were then multiplied by the per-cell ability of each phytoplankton strain to excrete fg of DOC per day and the fraction of each metabolite’s contribution to phytoplankton-excreted DOC. The total excretion flux was obtained by summing the values across all six phytoplankton classes. Note that each metabolite’s percentage contribution to phytoplankton-excreted DOC was extrapolated from a species to its corresponding phytoplankton class.

## Data availability

Raw mass spectrometry files have been deposited to the MetaboLights repository (https://www.ebi.ac.uk/metabolights/) and are accessible through https://www.ebi.ac.uk/metabolights/reviewer3182418e-6466-4e9a-8463-6844c8f1d021. Metadata exported from Skyline and MATLAB scripts used for metabolite data processing were uploaded to Github (https://github.com/yutingzhu0515/Characterization-of-Phytoplankton-Excreted-Metabolites). Metabolite concentration and cell counts data will be available at the designated BCO-DMO project page (https://www.bco-dmo.org/project/984095) upon publication.

## Acknowledgments

We acknowledge Noah Germolus and Brianna Garcia for their contributions to the MS data analysis pipeline development, Mary Ann Moran and Kathryn Halloran for contributions to data interpretation and discussion, and Laura Gray for data management. Funding is provided by the Center for Chemical Currencies of a Microbial Planet (C-CoMP) NSF-STC 2019589. This is C-CoMP publication 55.

## References

1. M. J. Behrenfeld, E. Boss, D. A. Siegel, D. M. Shea, Carbon-based ocean productivity and phytoplankton physiology from space. Global Biogeochemical Cycles 19 (2005).

2. M. A. Moran, et al., The Ocean’s labile DOC supply chain. Limnology and Oceanography 67, 1007–1021 (2022).

3. C. B. Field, M. J. Behrenfeld, J. T. Randerson, P. Falkowski, Primary Production of the Biosphere: Integrating Terrestrial and Oceanic Components. Science 281, 237–240 (1998).

4. G. E. Fogg, The Ecological Significance of Extracellular Products of Phytoplankton Photosynthesis. 26, 3–14 (1983).

5. B. Kieft, et al., Phytoplankton exudates and lysates support distinct microbial consortia with specialized metabolic and ecophysiological traits. Proceedings of the National Academy of Sciences 118, e2101178118 (2021).

6. M. A. Moran, et al., Microbial metabolites in the marine carbon cycle. Nat Microbiol 7, 508–523 (2022).

7. F. Azam, F. Malfatti, Microbial structuring of marine ecosystems. Nat Rev Microbiol 5, 782–791 (2007).

8. C. C. C. R. De Carvalho, P. Fernandes, Production of Metabolites as Bacterial Responses to the Marine Environment. Marine Drugs 8, 705–727 (2010).

9. C. Lønborg, C. Carreira, T. Jickells, X. A. Álvarez-Salgado, Impacts of Global Change on Ocean Dissolved Organic Carbon (DOC) Cycling. Frontiers in Marine Science 7 (2020).

10. B. P. Durham, et al., Recognition cascade and metabolite transfer in a marine bacteria-phytoplankton model system. Environmental Microbiology 19, 3500–3513 (2017).

11. F. X. Ferrer-González, et al., Resource partitioning of phytoplankton metabolites that support bacterial heterotrophy. The ISME journal 15, 762–773 (2021).

12. B. P. Durham, et al., An ecological framework for microbial metabolites in the ocean ecosystem. Limnology and Oceanography Letters 10, 636–659 (2025).

13. H. Alexander, et al., Functional group-specific traits drive phytoplankton dynamics in the oligotrophic ocean. Proceedings of the National Academy of Sciences 112, E5972– E5979 (2015).

14. Y. Sun, P. Debeljak, I. Obernosterer, Microbial iron and carbon metabolism as revealed by taxonomy-specific functional diversity in the Southern Ocean. ISME J 15, 2933–2946 (2021).

15. B. P. Durham, et al., Chemotaxonomic patterns in intracellular metabolites of marine microbial plankton. Frontiers in Marine Science 9, 864796 (2022).

16. J. Becker, et al., Closely related phytoplankton species produce similar suites of dissolved organic matter. Frontiers in Microbiology 5 (2014).

17. E. B. Kujawinski, et al., Metabolite diversity among representatives of divergent Prochlorococcus ecotypes. mSystems 8, e01261–22 (2023).

18. T. Anderson, H. Ducklow, Microbial loop carbon cycling in ocean environments studied using a simple steady-state model. Aquat. Microb. Ecol. 26, 37–49 (2001).

19. M. C. Kido Soule, K. Longnecker, W. M. Johnson, E. B. Kujawinski, Environmental metabolomics: Analytical strategies. Marine Chemistry 177, 374–387 (2015).

20. D. S. Wishart, et al., NMR and Metabolomics—A Roadmap for the Future. Metabolites 12, 678 (2022).

21. N. Zamboni, A. Saghatelian, G. J. Patti, Defining the Metabolome: Size, Flux, and Regulation. Molecular Cell 58, 699–706 (2015).

22. S. Pontrelli, U. Sauer, Salt-Tolerant Metabolomics for Exometabolomic Measurements of Marine Bacterial Isolates. Anal. Chem. 93, 7164–7171 (2021).

23. W. M. Johnson, M. C. Kido Soule, E. B. Kujawinski, Extraction efficiency and quantification of dissolved metabolites in targeted marine metabolomics. Limnology and Oceanography: Methods 15, 417–428 (2017).

24. J. S. Sacks, K. R. Heal, A. K. Boysen, L. T. Carlson, A. E. Ingalls, Quantification of dissolved metabolites in environmental samples through cation-exchange solid-phase extraction paired with liquid chromatography–mass spectrometry. Limnology and Oceanography: Methods 20, 683–700 (2022).

25. B. M. Garcia, et al., Benzoyl Chloride Derivatization Advances the Quantification of Dissolved Polar Metabolites on Coral Reefs. J. Proteome Res. 23, 2041–2053 (2024).

26. B. Widner, M. C. Kido Soule, F. X. Ferrer-González, M. A. Moran, E. B. Kujawinski, Quantification of Amine- and Alcohol-Containing Metabolites in Saline Samples Using Pre-extraction Benzoyl Chloride Derivatization and Ultrahigh Performance Liquid Chromatography Tandem Mass Spectrometry (UHPLC MS/MS). Anal. Chem. 93, 4809–4817 (2021).

27. R. Marcellin-Gros, G. Piganeau, D. Stien, Metabolomic Insights into Marine Phytoplankton Diversity. Mar Drugs 18, 78 (2020).

28. C. A. Carlson, S. Liu, B. M. Stephens, C. J. English, “Chapter 5 - DOM production, removal, and transformation processes in marine systems” in Biogeochemistry of Marine Dissolved Organic Matter (Third Edition), D. A. Hansell, C. A. Carlson, Eds. (Academic Press, 2024), pp. 137–246.

29. D. C. O. Thornton, Dissolved organic matter (DOM) release by phytoplankton in the contemporary and future ocean. European Journal of Phycology 49, 20–46 (2014).

30. J. W. Becker, S. L. Hogle, K. Rosendo, S. W. Chisholm, Co-culture and biogeography of Prochlorococcus and SAR11. ISME J 13, 1506–1519 (2019).

31. P. A. del Giorgio, J. J. Cole, Bacterial Growth Efficiency in Natural Aquatic Systems. Annual Review of Ecology and Systematics 29, 503–541 (1998).

32. S. Dutkiewicz, et al., Capturing optically important constituents and properties in a marine biogeochemical and ecosystem model. Biogeosciences 12, 4447–4481 (2015).

33. G. Forget, et al., ECCO version 4: an integrated framework for non-linear inverse modeling and global ocean state estimation. Geoscientific Model Development 8, 3071–3104 (2015).

34. B. A. Ward, S. Dutkiewicz, O. Jahn, M. J. Follows, A size-structured food-web model for the global ocean. Limnology and Oceanography 57, 1877–1891 (2012).

35. R. R. Malmstrom, M. T. Cottrell, H. Elifantz, D. L. Kirchman, Biomass Production and Assimilation of Dissolved Organic Matter by SAR11 Bacteria in the Northwest Atlantic Ocean. Applied and Environmental Microbiology 71, 2979–2986 (2005).

36. A. E. White, S. J. Giovannoni, Y. Zhao, K. Vergin, C. A. Carlson, Elemental content and stoichiometry of SAR11 chemoheterotrophic marine bacteria. Limnology and Oceanography Letters 4, 44–51 (2019).

37. H. Sarmento, et al., Phytoplankton species-specific release of dissolved free amino acids and their selective consumption by bacteria. Limnology and Oceanography 58, 1123–1135 (2013).

38. P. Peters, E. A. Galinski, H. G. Trüper, The biosynthesis of ectoine. FEMS Microbiology Letters 71, 157–162 (1990).

39. S. Fenizia, Thume, Wirgenings, Pohnert, Ectoine from Bacterial and Algal Origin Is a Compatible Solute in Microalgae. Marine Drugs 18, 42 (2020).

40. L. Czech, et al., Role of the Extremolytes Ectoine and Hydroxyectoine as Stress Protectants and Nutrients: Genetics, Phylogenomics, Biochemistry, and Structural Analysis. Genes (Basel) 9, 177 (2018).

41. C. L. Fiore, K. Longnecker, M. C. Kido Soule, E. B. Kujawinski, Release of ecologically relevant metabolites by the cyanobacterium Synechococcus elongatus CCMP 1631. Environmental Microbiology 17, 3949–3963 (2015).

42. T. M. D. Kasson, B. A. Barry, Reactive oxygen and oxidative stress: N-formyl kynurenine in photosystem II and non-photosynthetic proteins. Photosynth Res 114, 97–110 (2012).

43. C. P. Suffridge, et al., Exploring Vitamin B1 Cycling and Its Connections to the Microbial Community in the North Atlantic Ocean. Front. Mar. Sci. 7 (2020).

44. P. Carini, et al., Discovery of a SAR11 growth requirement for thiamin’s pyrimidine precursor and its distribution in the Sargasso Sea. The ISME Journal 8, 1727–1738 (2014).

45. M. Uchimiya, W. Schroer, M. Olofsson, A. S. Edison, M. A. Moran, Diel investments in metabolite production and consumption in a model microbial system. ISME J 16, 1306–1317 (2022).

46. W. F. Schroer, et al., Functional annotation and importance of marine bacterial transporters of plankton exometabolites. ISME COMMUN. 3, 1–10 (2023).

47. F. X. Ferrer-González, et al., Bacterial transcriptional response to labile exometabolites from photosynthetic picoeukaryote Micromonas commoda. ISME COMMUN. 3, 1–11 (2023).

48. B. P. Durham, et al., Sulfonate-based networks between eukaryotic phytoplankton and heterotrophic bacteria in the surface ocean. Nature microbiology 4, 1706–1715 (2019).

49. F. Götz, et al., Targeted metabolomics reveals proline as a major osmolyte in the chemolithoautotroph Sulfurimonas denitrificans. MicrobiologyOpen 7, e00586 (2018).

50. D. D. Martin, R. A. Ciulla, M. F. Roberts, Osmoadaptation in Archaea. Applied and Environmental Microbiology 65, 1815–1825 (1999).

51. H. Ono, et al., Characterization of biosynthetic enzymes for ectoine as a compatible solute in a moderately halophilic eubacterium, Halomonas elongata. J Bacteriol 181, 91–99 (1999).

52. A. Schulz, et al., Feeding on compatible solutes: A substrate-induced pathway for uptake and catabolism of ectoines and its genetic control by EnuR. Environmental Microbiology 19, 926–946 (2017).

53. R. R. L. Guillard, P. E. Hargraves, Stichochrysis immobilis is a diatom, not a chrysophyte. Phycologia 32, 234–236 (1993).

54. J. B. Waterbury, S. W. Watson, F. W. Valois, D. G. Franks, “Biological and ecological characterization of the marine unicellular cyanobacterium Synechococcus” in Platt, T., Li, W.K.I. Photosynthetic Picoplankton, (Can. Bull. Fish. Aquatic. Sci, 1986), p. 214.

55. L. R. Moore, et al., Culturing the marine cyanobacterium Prochlorococcus. Limnology and Oceanography: Methods 5, 353–362 (2007).

56. K. J. Adams, et al., Skyline for Small Molecules: A Unifying Software Package for Quantitative Metabolomics. J Proteome Res 19, 1447–1458 (2020).

57. C. M. Henderson, N. J. Shulman, B. MacLean, M. J. MacCoss, A. N. Hoofnagle, Skyline Performs as Well as Vendor Software in the Quantitative Analysis of Serum 25-Hydroxy Vitamin D and Vitamin D Binding Globulin. Clin Chem 64, 408–410 (2018).

58. Zar, Biostatistical Analysis, 4th Ed. (Prentice Hall, 1999).

59. Y. Djoumbou Feunang, et al., ClassyFire: automated chemical classification with a comprehensive, computable taxonomy. Journal of Cheminformatics 8, 61 (2016).

60. E. V. Armbrust, et al., The Genome of the Diatom Thalassiosira Pseudonana: Ecology, Evolution, and Metabolism. Science 306, 79–86 (2004).

61. S. R. Bench, et al., Whole genome comparison of six Crocosphaera watsonii strains with differing phenotypes. Journal of Phycology 49, 786–801 (2013).

62. G. C. Kettler, et al., Patterns and Implications of Gene Gain and Loss in the Evolution of Prochlorococcus. PLOS Genetics 3, e231 (2007).

63. B. Palenik, et al., The genome of a motile marine Synechococcus. Nature 424, 1037–1042 (2003).

64. A. Z. Worden, et al., Green Evolution and Dynamic Adaptations Revealed by Genomes of the Marine Picoeukaryotes Micromonas. Science 324, 268–272 (2009).

65. O. Nam, J.-M. Park, H. Lee, E. Jin, De novo transcriptome profile of coccolithophorid alga Emiliania huxleyi CCMP371 at different calcium concentrations with proteome analysis. PLoS ONE 14, e0221938 (2019).

66. M. Kanehisa, Y. Sato, M. Kawashima, M. Furumichi, M. Tanabe, KEGG as a reference resource for gene and protein annotation. Nucleic Acids Res 44, D457–D462 (2016).

67. A. M. Eren, et al., Community-led, integrated, reproducible multi-omics with anvi’o. Nat Microbiol 6, 3–6 (2021).

68. M. D. Ashkezari, et al., Simons Collaborative Marine Atlas Project (Simons CMAP): An open-source portal to share, visualize, and analyze ocean data. Limnology and Oceanography: Methods 19, 488–496 (2021).

69. S. Bertilsson, O. Berglund, D. M. Karl, S. W. Chisholm, Elemental composition of marine Prochlorococcus and Synechococcus: Implications for the ecological stoichiometry of the sea. Limnology and Oceanography 48, 1721–1731 (2003).

70. S. Menden-Deuer, E. J. Lessard, Carbon to volume relationships for dinoflagellates, diatoms, and other protist plankton. Limnology and Oceanography 45, 569–579 (2000).

71. W.-D. Lu, Z.-M. Chi, C.-D. Su, Identification of glycine betaine as compatible solute in Synechococcus sp. WH8102 and characterization of its N-methyltransferase genes involved in betaine synthesis. Arch Microbiol 186, 495–506 (2006).

