## Supplemental Information for "Characterization of Phytoplankton-Excreted Metabolites Mediating Carbon Flux through the Surface Ocean"

**Table S1.** Lists of names and abbreviations of metabolites with the chosen SIL-IS types and polarities, along with limits of detection (LOD) and limits of linearity (LOL). The concentrations of 2′-deoxyguanosine, 3′-deoxyguanosine, ciliatine, cytosine, folic acid, muramic acid, and tryptamine were either not above the LOD or were not significantly different from the media blanks and these metabolites were excluded from the list of detected exometabolites.

| Compound | Abbreviation | Polarity | SIL-IS | LOD (nM) | LOL (nM) |
|---|---|---|---|---|---|
| 2′-deoxyguanosine [1] | | positive | $D_5$ | 2.3E+00 | 748 |
| 2′-deoxyuridine [1] | | negative | $D_5$ | 7.2E-01 | 876 |
| adenosine 3′-monophosphate [1] | 3′AMP | positive | $D_5$ | 1.6E-02 | 432 |
| 3′-deoxyguanosine [1] | | positive | $D_5$ | 2.1E-01 | 748 |
| adenosine 5′-monophosphate [1] | 5′AMP | positive | $D_5$ | 1.4E-01 | 511 |
| inosine 5′-monophosphate [1] | 5′IMP | negative | $D_5$ | 1.1E-01 | 255 |
| uridine 5′-monophosphate [1] | 5′UMP | negative | $D_5$ | 6.0E-02 | 272 |
| 2,3-dihydroxypropane-1-sulfonate [1] | DHPS | negative | $D_5$ | 2.4E-02 | 1281 |
| γ-aminobutyric acid [1] | GABA | negative | $D_5$ | 2.5E-02 | 2909 |
| 4-amino-2-methyl-5-pyrimidinemethanol [1] | HMP | positive | $D_5$ | 4.7E-02 | 1078 |
| N-acetyl-muramic acid | | positive | $D_5$ | 1.0E+00 | 511 |
| S-(1,2-dicarboxyethyl)glutathione | | positive | $D_5$ | 1.0E-02 | 709 |
| adenosine [1] | | positive | $D_5$ | 3.3E-01 | 374 |
| alanine [1] | | negative | $^{13}C_6$ | 3.9E-01 | 2245 |
| 4-amino-5-aminomethyl-2-methylpyrimidine [1] | amMP | negative | $D_5$ | 1.7E+00 | 1086 |
| arginine [1] | | positive | $^{13}C_6$ | 2.3E-01 | 1722 |
| asparagine [1] | | positive | $^{13}C_6$ | 1.1E-01 | 757 |
| aspartic acid [1] | | negative | $^{13}C_6$ | 3.4E-01 | 2254 |
| chitobiose [1] | | positive | $D_5$ | 1.6E-01 | 353 |
| chitotriose [1] | | positive | $^{13}C_6$ | 2.1E-02 | 478 |
| ciliatine [1] | | positive | $D_5$ | 6.4E-03 | 2399 |
| citrulline [1] | | negative | $D_5$ | 2.9E-01 | 1712 |
| cysteic acid [1] | | positive | $D_5$ | 4.3E-02 | 1774 |
| cysteine [2] | | positive | $D_5$ | 4.5E-01 | 2476 |
| cytidine [1] | | positive | $D_5$ | 3.7E-02 | 822 |
| cytosine [1] | | positive | $D_5$ | 2.3E-01 | 2700 |
| ectoine [1] | | positive | $D_5$ | 4.1E-01 | 2110 |
| folic acid [1] | | positive | $D_5$ | 1.2E-01 | 340 |
| glucosamine-6-phosphate [1] | | negative | $D_5$ | 6.6E-02 | 658 |
| glucose 6-phosphate [1] | | negative | $D_5$ | 5.7E-02 | 132 |
| glutamic acid [1 *] | | positive | $^{13}C_6$ | 2.2E-01 | 2039 |
| glutamine [1] | | positive | $^{13}C_6$ | 1.6E-01 | 1026 |

| | | | | |
|---|---|---|---|---|
| glutathione [2] | positive | $^{13}C_6$ | 5.8E-03 | 325 |
| glycine [1] | positive | $D_5$ | 1.2E+00 | 3996 |
| guanosine [1] | positive | $D_5$ | 4.3E-02 | 1059 |
| histidine [1] | negative | $^{13}C_6$ | 2.3E-01 | 1934 |
| homoserine [1] | positive | $^{13}C_6$ | 8.5E-02 | 1259 |
| inosine [1] | positive | $D_5$ | 3.8E-01 | 373 |
| isethionic acid [1] | negative | $D_5$ | 2.0E-02 | 675 |
| isoleucine [1] | negative | $^{13}C_6$ | 6.0E-02 | 2287 |
| kynurenine [1] | negative | $D_5$ | 4.1E-02 | 1441 |
| leucine [1] | positive | $D_5$ | 1.7E-01 | 1144 |
| lysine [2] | negative | $^{13}C_6$ | 1.8E-01 | 1368 |
| malic acid [1] | negative | $D_5$ | 1.6E+00 | 1119 |
| methionine [1] | negative | $D_5$ | 3.1E+00 | 2011 |
| muramic acid [2] | negative | $^{13}C_6$ | 1.1E-01 | 796 |
| ornithine [2] | negative | $^{13}C_6$ | 1.7E-01 | 1776 |
| pantothenic acid [1] | positive | $D_5$ | 1.9E-02 | 630 |
| phenylalanine [1] | negative | $^{13}C_6$ | 3.6E-02 | 1211 |
| proline [1] | negative | $^{13}C_6$ | 3.4E-01 | 2606 |
| putrescine [2] | negative | $^{13}C_6$ | 2.9E-02 | 1242 |
| sarcosine [1] | positive | $D_5$ | 7.4E-02 | 3367 |
| serine [1] | positive | $D_5$ | 1.5E+00 | 1427 |
| glycerol 3-phosphate [1] | negative | $D_5$ | 5.1E-02 | 540 |
| spermidine [3] | negative | $^{13}C_6$ | 2.2E-02 | 688 |
| taurine [1] | positive | $D_5$ | 4.0E-02 | 2397 |
| threonine [1] | negative | $D_5$ | 2.4E-01 | 2518 |
| tryptamine [1] | positive | $D_5$ | 1.0E-01 | 1872 |
| tryptophan [1] | negative | $D_5$ | 6.6E-02 | 734 |
| tyrosine [1] | negative | $D_5$ | 8.3E-01 | 552 |
| uridine [1] | negative | $D_5$ | 5.0E-02 | 819 |
| valine [1] | negative | $^{13}C_6$ | 2.1E-01 | 2561 |
| xanthosine [1] | positive | $D_5$ | 3.4E-01 | 624 |

[1,2,3] The number of benzoyl labels on the compound

[*] O-acetyl-L-serine is an interfering peak for glutamic acid but was not originally included in our targeted list for this work's metabolite analysis. Therefore glutamic acid concentrations presented here are upper limit values. O-acetyl-L-serine concentrations in filtrate samples of *C. watsonii* and *M. commoda* were analyzed separately using an aniline derivatization method (Halloran et al., in prep) and were below the detection limits for both species.

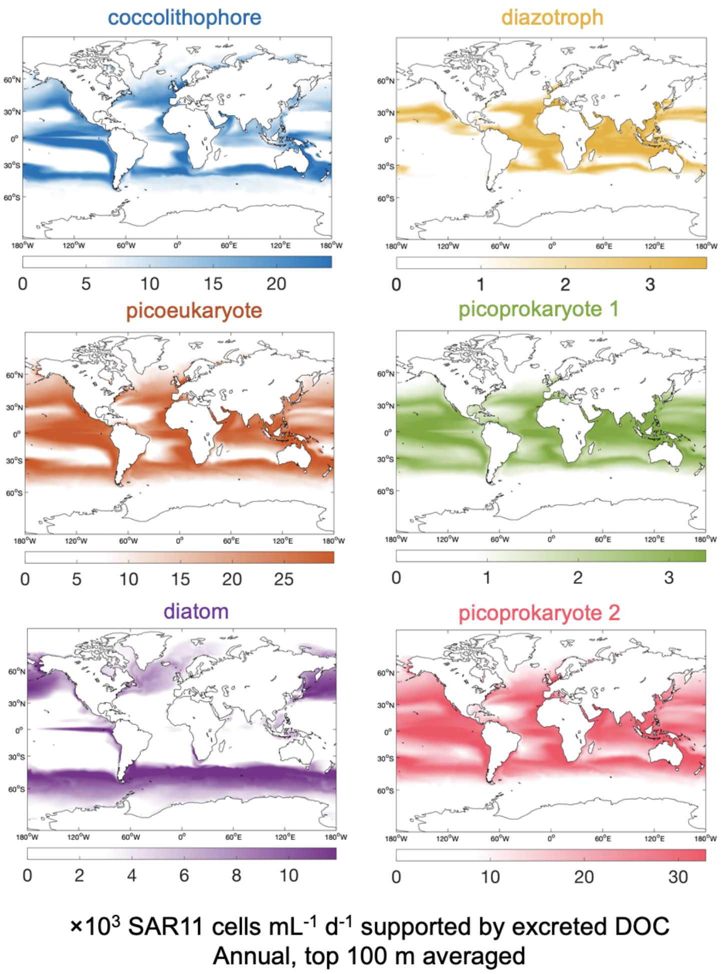


**Figure S1.** Global distribution of computed numbers of new bacterial cells that can be produced from total DOC excreted by coccolithophores, picoeukaryotes, diatoms (< 15 µm fractions were selected), diazotrophs, and picoprokaryote size fractions 1 (analogue for *Prochlorococcus*) and 2 (analogue for *Synechococcus*), in per mL of seawater per day. 54% BGE (1, 2) were assumed. Phytoplankton biomass climatology (in mmol C/m$^3$) output data of the MIT Darwin model were downloaded from Simons Collaborative Marine Atlas Project (https://simonscmap.com/).

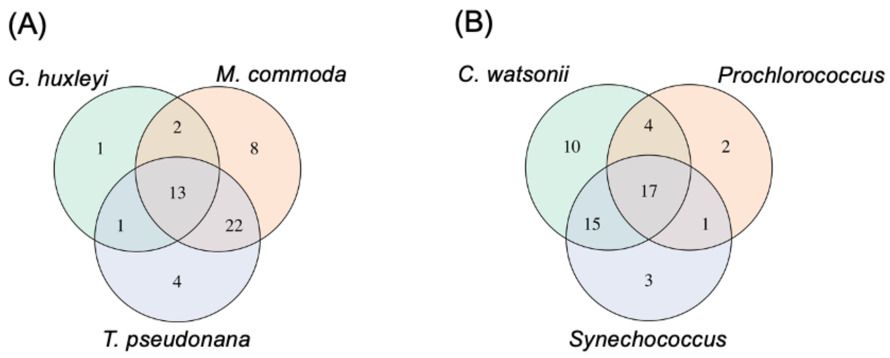


**Figure S2.** Venn diagrams representing shared exometabolites among (A) *G. huxleyi*, *M. commoda*, and *T. pseudonana* and (B) *C. watsonii, Synechococcus,* and *Prochlorococcus*.

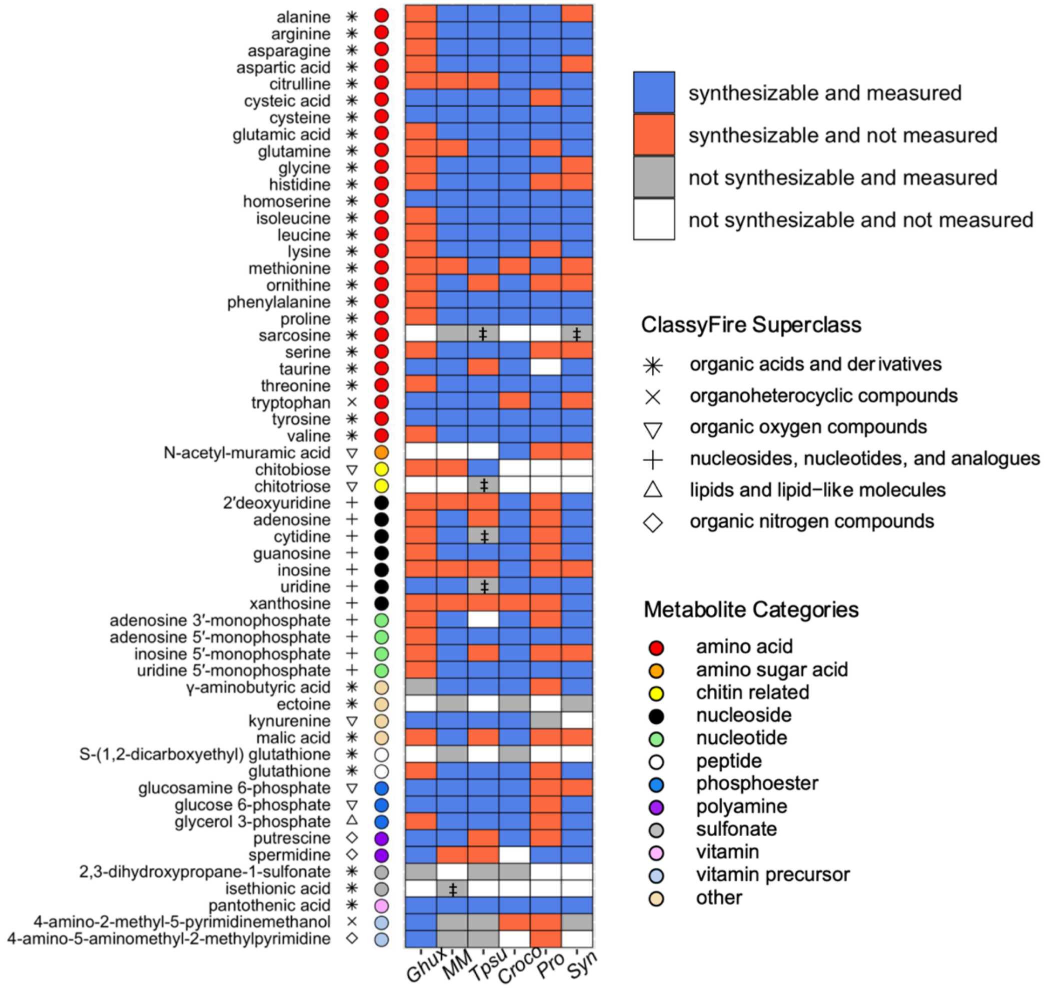


**Figure S3.** Comparison of biosynthesizable exometabolites and the measured pool of exometabolites in axenic cultures of *G. huxleyi* (*Ghux*), *C. watsonii* (*Croco*), *M. commoda* (*MM*), *Prochlorococcus* (*Pro*), *Synechococcus* (*Syn*), and *T. pseudonana* (*Tpsu*). Each identified exometabolite is paired with a symbol representing ClassyFire (3) superclass and a colored marker that represents the assigned metabolite category in this work. Metabolites were ordered according to their assigned metabolite categories. For metabolites marked with a double dagger symbol (‡), they should be synthesizable despite not being found in KEGG pathways based on known metabolite functionality or homologous enzymes.

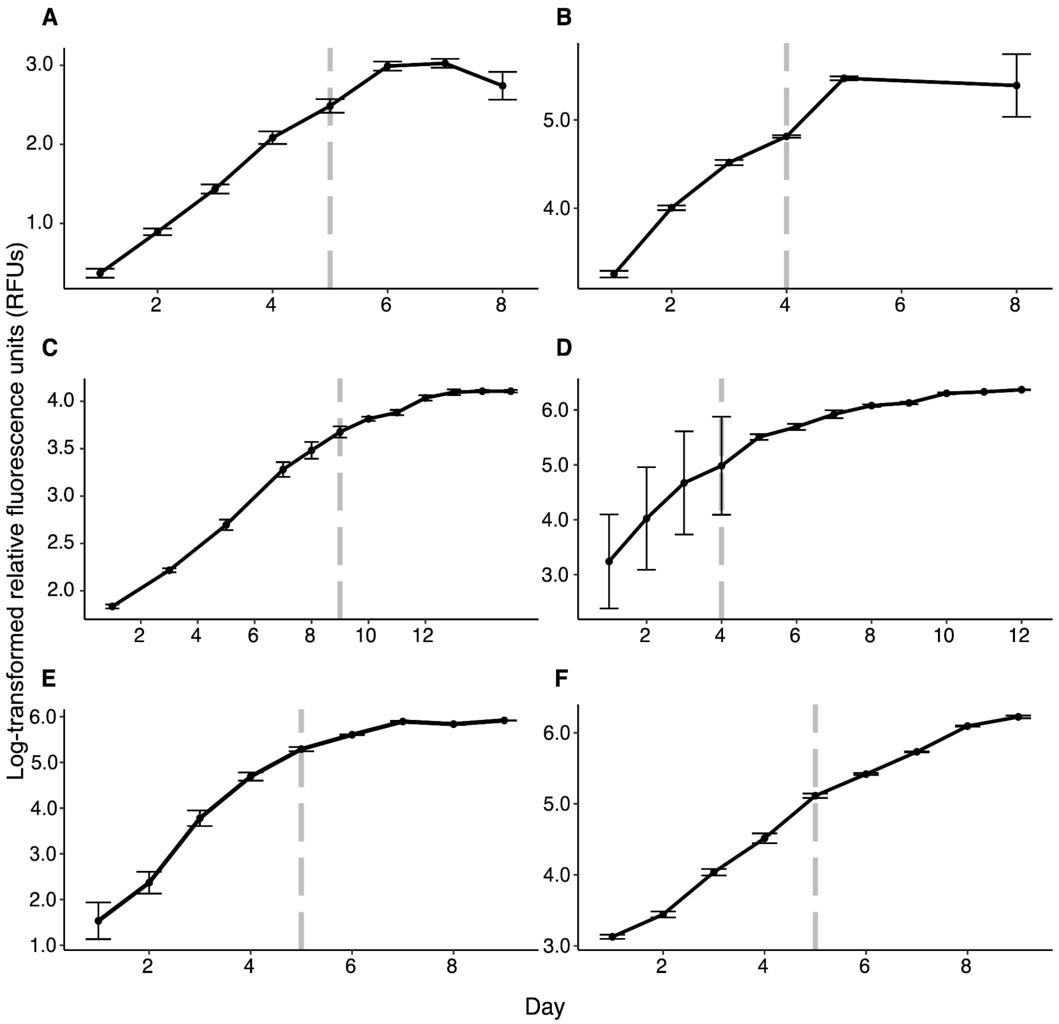


**Figure S4.** Growth curves of (A) *Synechococcus* sp. WH8102, (B) *Prochlorococcus marinus* MIT 9301, (C) *Crocosphaera watsonii* WH8501, (D) *Thalassiosira pseudonana* CCMP1335, (E) *Gephyrocapsa huxleyi* CCMP371, and (F) *Micromonas commoda* RCC299 measured by log-transformed relative fluorescence units (RFUs). Log-transformed RFUs were averaged across nine biological replicates until six replicates were harvested, after which three replicates were monitored for all taxa except *Prochlorococcus* where two replicates were monitored. Vertical grey dashed lines represent the date of harvest in each taxa; log-transformed RFUs after this date were averaged across the remaining replicates. Error bars represent standard deviations calculated from log-transformed RFUs.

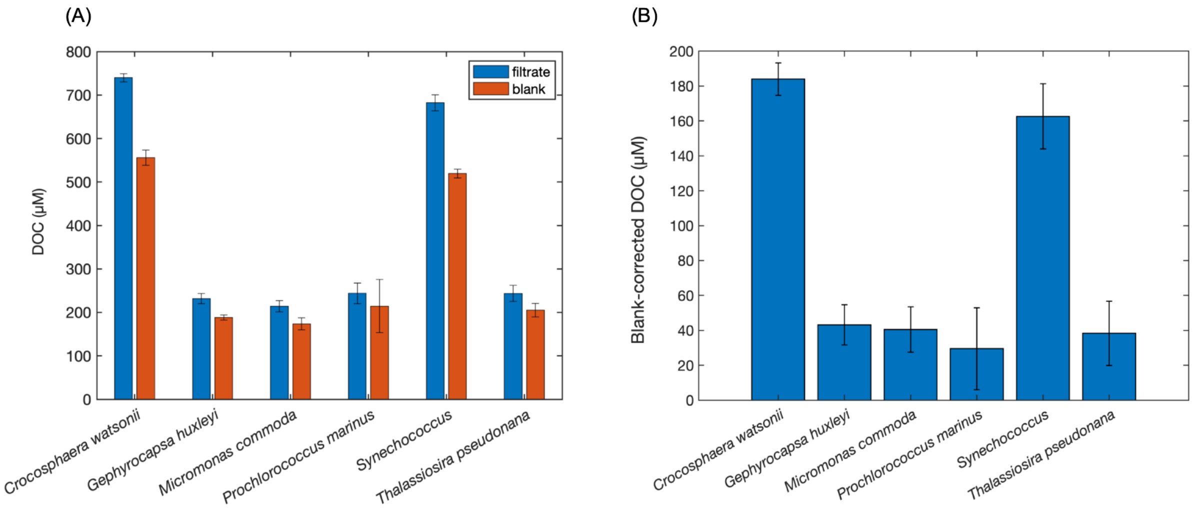


**Figure S5.** (A) Average DOC concentrations in filtrate samples (N = 6) and media blanks (N = 2 – 4), and (B) average blank-corrected DOC concentrations in culture samples of *C. watsonii*, *G. huxleyi*, *M. commoda*, *Synechococcus*, *Prochlorococcus*, and *T. pseudonana*. Error bars denote standard deviations (N=6).

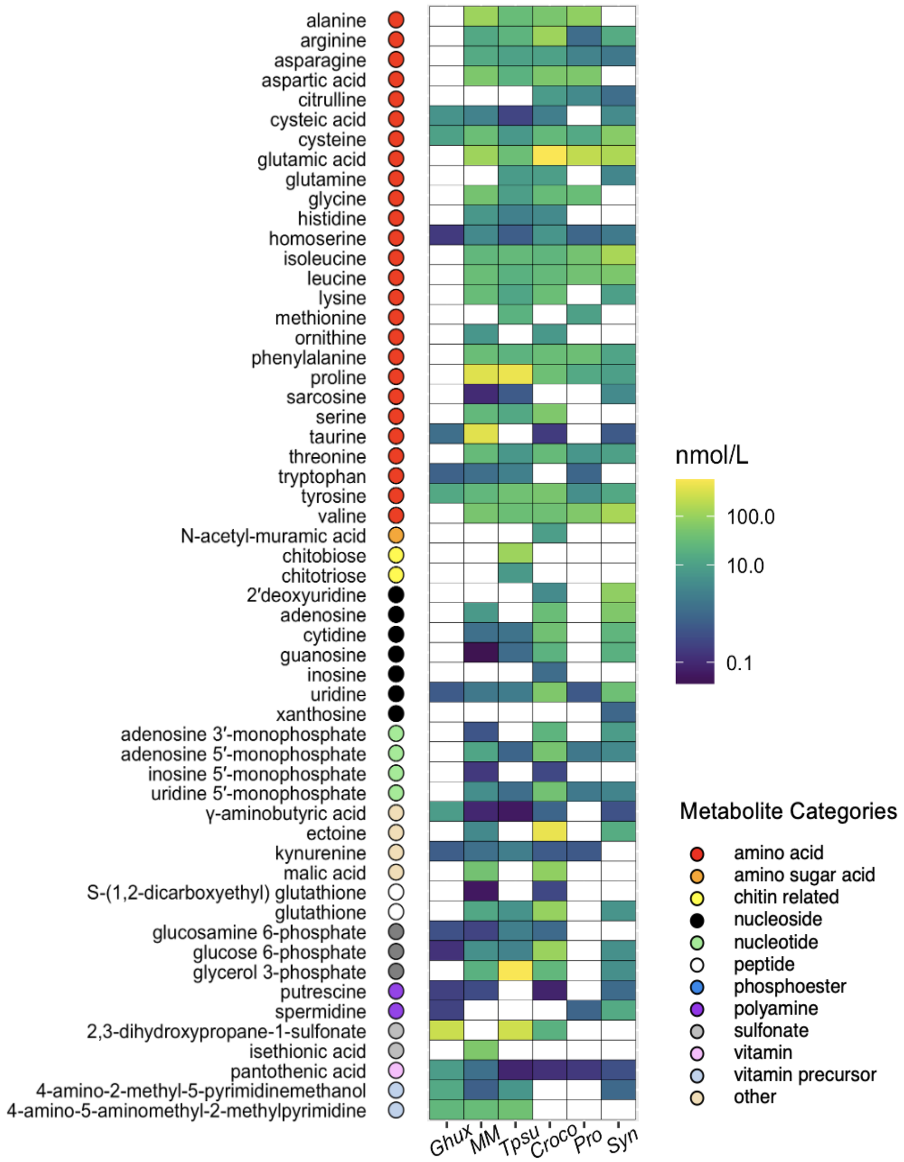


**Figure S6.** Nanomolar exometabolite concentrations in axenic cultures of *C. watsonii* (*Croco*), *G. huxleyi* (*Ghux*), *M. commoda* (*MM*), *Prochlorococcus* (*Pro*), *Synechococcus* (*Syn*), and *T. pseudonana* (*Tpsu*). Nanomolar concentrations were obtained by subtracting average concentrations in filtrate samples (N = 6) by average concentrations in media blanks (N = 2–4). Each identified exometabolite is paired with a colored marker that represents the assigned metabolite category.

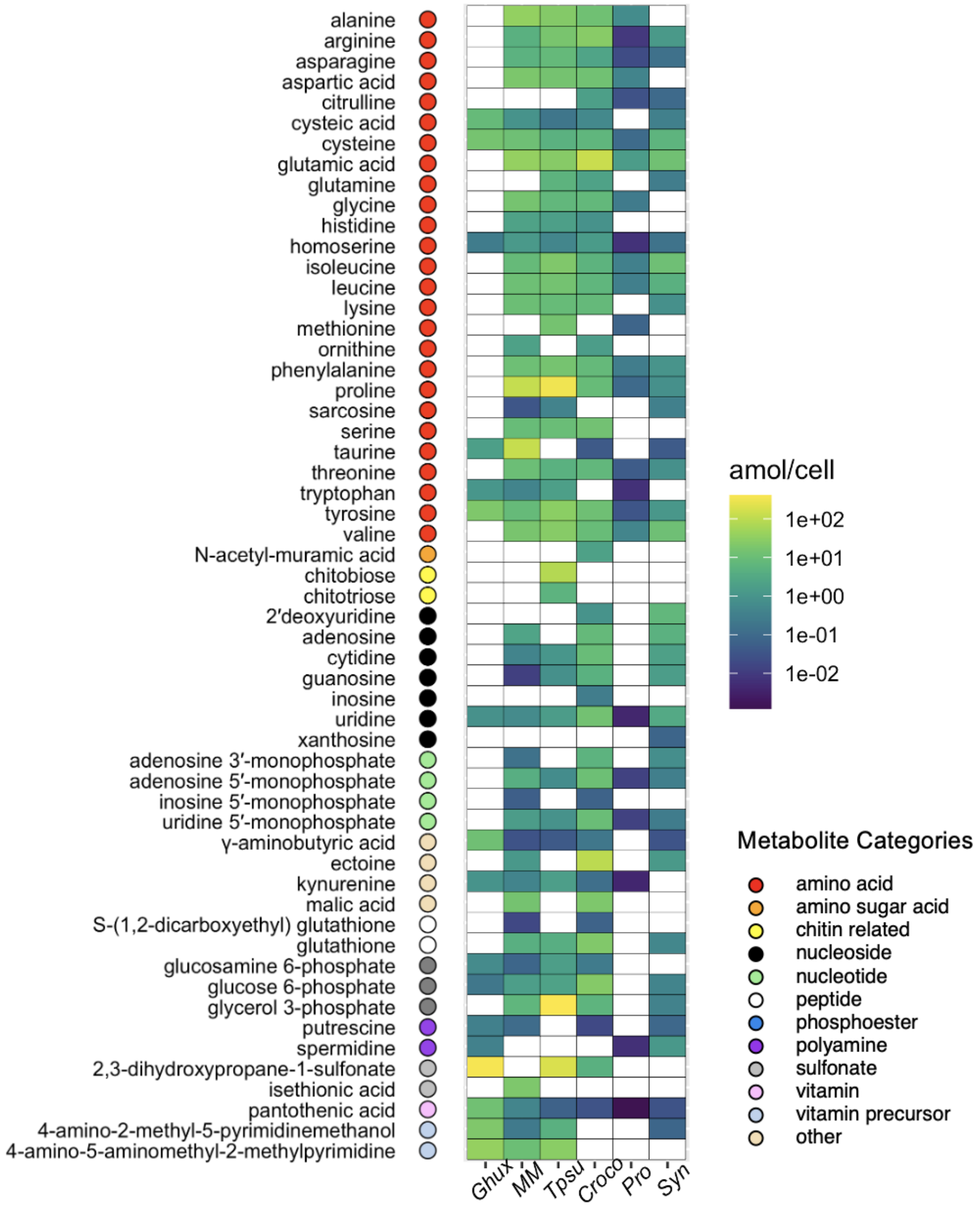


**Figure S7.** Cell-specific concentrations of exometabolite in axenic cultures of *C. watsonii* (*Croco*), *G. huxleyi* (*Ghux*), *M. commoda* (*MM*), *Prochlorococcus* (*Pro*), *Synechococcus* (*Syn*), and *T. pseudonana* (*Tpsu*).

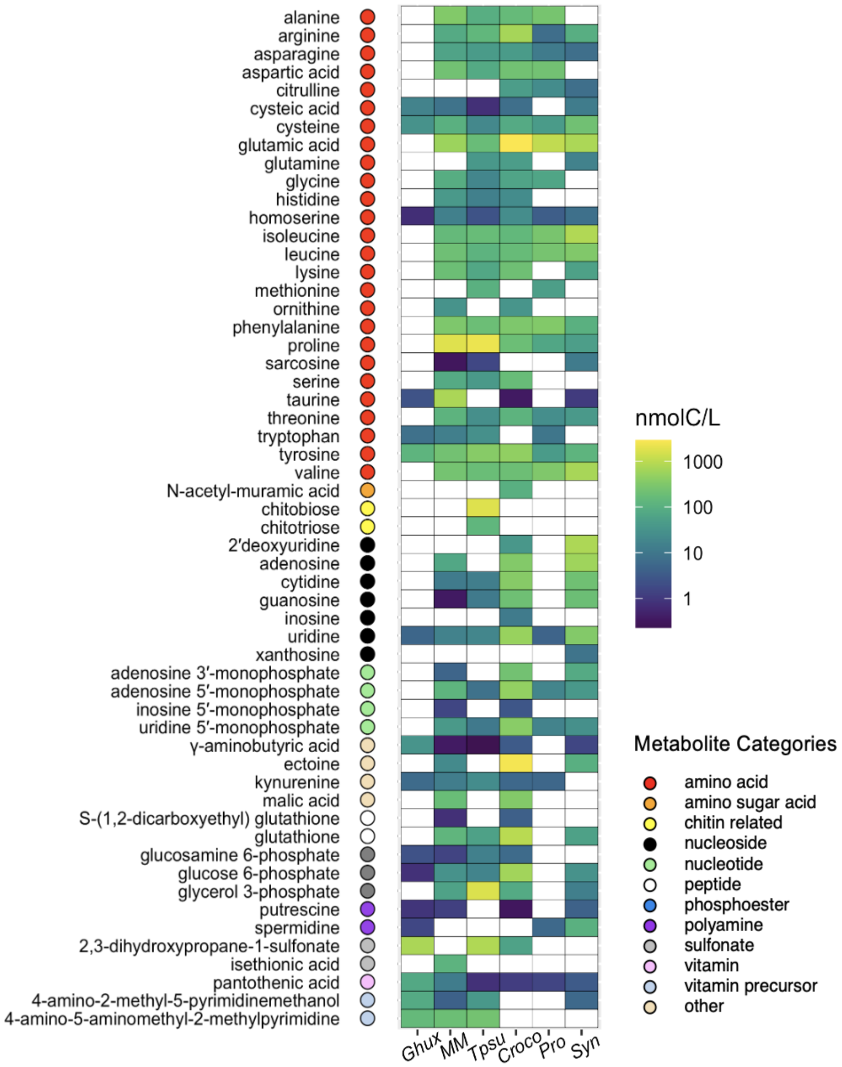


**Figure S8.** Carbon concentrations of exometabolite in axenic cultures of *C. watsonii* (*Croco*), *G. huxleyi* (*Ghux*), *M. commoda* (*MM*), *Prochlorococcus* (*Pro*), *Synechococcus* (*Syn*), and *T. pseudonana* (*Tpsu*). Carbon concentrations were obtained by multiplying blank-corrected, average nanomolar concentrations by number of carbon atoms in each molecule. Each identified exometabolite is paired with a colored marker that represents the assigned metabolite category.

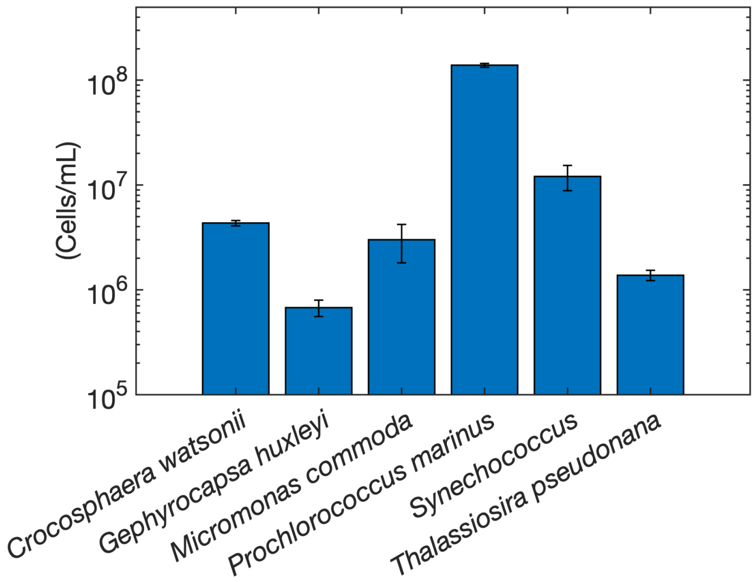


**Figure S9.** Average cell counts in culture samples of *C. watsonii*, *G. huxleyi*, *M. commoda*, *Prochlorococcus*, *Synechococcus*, and *T. pseudonana*. Error bars denote standard deviations (N=6).

### Supplementary Text

Isethionic acid, an organic sulfur compound, was detected in *M. commoda*, which is consistent with its previous detection in the endometabolome of *M. pusilla* (4). Isethionic acid has no KEGG-annotated biosynthetic pathway in *M. commoda* (Figure S3), however *M. pusilla* encodes homologs of the two proteins, taurine aminotransferase (*tpa*/*toa*) and sulfoacetaldehyde reductase (*isfD*), required for isethionate production from taurine in the characterized bacterial catabolic pathway (5). Homologs of these enzymes are found in the *M. commoda* genome (81% amino acid identity with putative *M. pusilla tpa*, 84% amino acid identity with putative *M. pusilla isfD*), indicating a putative biosynthetic pathway for isethionic acid could have been produced in the exometabolome.

Chitin-related exometabolite chitotriose was detected and did not have a KEGG-annotated biosynthetic pathway in *T. pseudonana* (Figure S3). However, the *T. pseudonana* genome encodes several chitinases, and endochitinases are capable of depolymerizing chitin to chitobiose and other oligomers like chitotriose and chitotetrose, so a biosynthesis pathway for chitotriose in *T. pseudonana* likely exists. In these and other related examples, the lack of an annotated biosynthetic pathway for a detected exometabolite is likely a function of inaccurate gene calling or incomplete pathway annotation.

Sarcosine was detected as an exometabolite for *M. commoda*, *T. pseudonana*, and *Synechococcus*, without annotated biosynthesis pathways in KEGG (Figure S3). However, two homologous genes coding for glycine (sarcosine *N*-methyltransferase and sarcosine dimethylglycine *N*-methyltransferase, which regulate glycine to sarcosine biosynthesis and sarcosine to betaine biosynthesis, respectively) were identified by Lu et al. (6) for *Synechococcus* WH8102. In addition, the *T. pseudonana* CCMP1335 genome encodes sarcosine dehydrogenase (Gene ID: 7444857), which catalyzes the oxidation of sarcosine to glycine. These findings highlight both the general issue of missing or incorrect annotations that can occur within genomic repositories, and the potential for metabolomics in improving annotations in species of interest.

Cytidine and uridine were detected as exometabolites in *T. pseudonana*, yet KEGG does not annotate a complete biosynthetic pathway for pyrimidine nucleotides in this species (Figure S3). However, the *T. pseudonana* CCMP1335 genome encodes CTP synthase (Gene ID 7444132), which interconverts UTP and CTP. In addition, the genome contains genes involved in pyrimidine salvage and interconversion, including uracil phosphoribosyltransferase (Gene ID 7445346), pyrimidine-nucleoside phosphorylase (Gene ID 7445357), and 5'-nucleotidase (Gene ID 7445357), which together mediate nucleotide dephosphorylation and recycle uracil, uridine, and cytidine into nucleotide pools. Gene IDs refer to NCBI Gene entries for *T. pseudonana* CCMP1335 (https://www.ncbi.nlm.nih.gov/datasets/gene/GCF_000149405.2/; accessed October 2025).

### References


1. M. A. Moran, et al., The Ocean's labile DOC supply chain. Limnology and Oceanography 67, 1007–1021 (2022).
2. P. A. del Giorgio, J. J. Cole, Bacterial Growth Efficiency in Natural Aquatic Systems. Annual Review of Ecology and Systematics 29, 503–541 (1998).
3. Y. Djoumbou Feunang, et al., ClassyFire: automated chemical classification with a comprehensive, computable taxonomy. Journal of Cheminformatics 8, 61 (2016).
4. B. P. Durham, et al., Chemotaxonomic patterns in intracellular metabolites of marine microbial plankton. Frontiers in Marine Science 9, 864796 (2022).
5. B. P. Durham, et al., Sulfonate-based networks between eukaryotic phytoplankton and heterotrophic bacteria in the surface ocean. Nature microbiology 4, 1706–1715 (2019).
6. W.-D. Lu, Z.-M. Chi, C.-D. Su, Identification of glycine betaine as compatible solute in Synechococcus sp. WH8102 and characterization of its N-methyltransferase genes involved in betaine synthesis. Arch Microbiol 186, 495–506 (2006).
